# Matrix Stiffness-driven FAK Splicing Tunes Cell Mechanosensing

**DOI:** 10.64898/2026.07.25.736411

**Authors:** Martial Millet, Juliette Gouhier, Olivier Chancy, Elise Tahon, William David Zakzuk Vivas, Emilie Pic, Gabriel Khelifi, Solange Landreville, Samer Hussein, Dwayne Stupack, François Bordeleau

**Affiliations:** Centre de recherche du CHU de Québec-Université Laval, Québec, QC, Canada; Université Laval Cancer Research Center, Quebec City, QC, Canada; Université Laval Tissue Engineering Laboratory (LOEX) Research Center, Quebec City, QC, Canada; Department of Ophthalmology and Otorhinolaryngology-Cervico-Facial Surgery, Faculty of Medicine, Université Laval, Quebec City, QC, Canada; Department of Molecular Biology, Medical Biochemistry and Pathology, Faculty of Medicine, Université Laval, Quebec City, QC, Canada; Department of Obstetrics, Gynecology, and Reproductive Sciences, Moores Cancer Center, Division of Gynecologic Oncology, University of California San Diego, La Jolla, CA, USA

**Keywords:** Alternative Splicing, Cell Migration, Durotaxis, ECM Stiffness, Focal Adhesion Kinase, Mechanosensing, Mechanotransduction

## Abstract

The extracellular matrix (ECM) stiffness influences many physiological and pathological processes. Cells can sense mechanical changes within the ECM through mechanosensors, most notably through the focal adhesion kinase (FAK). Using human-derived data and 2D/3D engineered models, we identified a ubiquitous and uncharacterized FAK isoform lacking exon 4 (FAKΔe4). We demonstrated that FAKΔe4 splicing is regulated by substrate stiffness in a biphasic manner, in which an optimal stiffness is required for a maximal FAKΔe4 expression. We further showed that FAKΔe4 dictates at which stiffness optimal migration speed and invasion occur, impairs focal adhesion dynamics and maturation, and shifts its autophosphorylation and downstream YAP nuclear translocation toward lower stiffness compared to the canonical FAK isoform. Moreover, the FAKΔe4 expression levels to canonical FAK determine at which stiffness cells will converge during durotaxis. Our results reveal how FAKΔe4 acts as a fine-tuning mechanosensing switch and reframe our understanding of mechanotransduction.

## Introduction

The extracellular matrix (ECM) stiffness is essential to maintain tissue homeostasis. Alterations of the ECM mechanical properties can drive both developmental and pathological processes (1,2). For instance, ECM stiffening is now appreciated as a major driver of tumor progression toward a metastatic disease (3,4). Increased ECM stiffness promotes biological processes such as cell differentiation, spreading, migration, invasion and metastasis (5–8). One of the main structures by which cells can sense the stiffness of their surrounding matrix is through focal adhesions (FAs) (9). FAs are dynamic multiprotein complexes that act both as the mechanical coupler between the ECM and the cytoskeleton and as the integrator to turn those mechanical cues into biochemical cues (9,10). Interestingly, depending on their invasive phenotype, tumor cells will not respond in the same way to similar ECM stiffness-related cues. Notably, cell migration displays a biphasic relationship relative to ECM stiffness, and the optimal migration speed of invasive cells can occur at higher stiffness than less invasive cells (11,12). Moreover, when presented with a stiffness gradient, cells are able to identify an optimal stiffness to which they will converge (13–16). The optimal stiffness for a tumor cell appears to be linked to its aggressiveness (17). In addition, downregulating FA structural proteins can shift the optimal stiffness that a cell will prefer (11,12,18,19). However, while formation and dynamics of FAs appear central to how cells determine their optimal ECM stiffness, the molecular mechanosensors underlying this process remain largely unknown.

The main mechanosensing protein in FAs is the non-receptor tyrosine Focal Adhesion Kinase (FAK, also named PTK2). High ECM stiffness activates integrin signaling through force transduction, leading to the activation of FAK by triggering its autophosphorylation on Y397 (9). Specifically, force-induced FAK unfolding promotes an open conformation that enables this autophosphorylation (20). FAK autophosphorylation is a key regulatory step to initiate downstream stiffness-mediated signaling that controls FA dynamics and maturation as well as subsequent cell migration (21–23). Stiffness-dependent FAK activation is required to trigger RhoA-dependent contractility and mechano-dependent YAP nuclear translocation (9,24). In turn, increased cell contractility is critical to FA maturation (25). Conversely, FAK is also involved in controlling FA disassembly dynamic (23). Importantly, FAK was shown to influence durotaxis events, suggesting it could act as a molecular switch that allows cells to determine their optimal ECM stiffness (26).

Alternative splicing is a fundamental mechanism that allows the expression of different protein isoforms, thereby increasing protein diversity (27–29). Small expression changes in alternatively spliced isoforms can modulate cell behaviors and phenotypes (30,31). However, solid tumors display an aberrant alternative splicing profile compared to healthy tissue favoring tumor development (28,32). Both hallmarks of cancer, alternative splicing and ECM stiffness have been linked together. Indeed, previous work showed that ECM stiffness regulates alternative splicing events (30,33). Interestingly, stiffness-driven splicing events can alter tumor cell response as a function of the underlaying stiffness, including stiffness–induced tumor cell extravasation and proliferation (30,34). On the other hand, the alternative splicing in talin influences its force-dependent unfolding, which impacts cell adhesion and motility (35), further highlighting a role for splicing in regulating mechanosensing. Interestingly, several FAK splice variants have been identified in tumor samples, including ovary and colorectal, and were shown to display increased basal autophosphorylation levels (36–38). However, whether ECM stiffness regulates FAK alternative splicing, which in turn would influence tumor cell behavior through altered mechanosensing, remains uncharted.

Here, we identified and explored the functions of an uncharacterized stiffness-regulated FAK splice isoform, FAKΔe4, which we found to be ubiquitously expressed across all tissue types and upregulated in various cancers. Using both 2D and 3D engineered substrates, we showed that FAKΔe4 displays altered mechano-induced autophosphorylation, disrupts FA dynamics and maturation, thereby modulating mechanosensing and cell migration in a stiffness-dependent manner. Mechanistically, FAKΔe4 alters YAP-dependent mechanotransduction and tunes the optimal ECM stiffness sensed by tumor cells during durotaxis. Together, our findings identify FAKΔe4 as a previously unrecognized regulator of cell mechanosensing.

## Results

### FAKΔe4 is expressed across normal and tumor tissues

Given the key role of FAK in cancer progression and the impact of alternative splicing on protein function (39,40), we investigated whether the alternative splicing of FAK is altered in tumors. We first quantified exon inclusion quantified as the percent spliced-in (PSI; e.g. a lower PSI means a higher expression of the FAKΔe4) for the FAK splicing events previously reported in the literature, namely the inclusions of either exon 14, 16 or 31 (36–38). Considering seven different target human tissues, we found that those three specific splicing events were rarely differentially expressed between normal and the corresponding tumor tissues (Fig.S1A-C). However, we identified an uncharacterized FAK splicing event that corresponds to the exclusion of exon 4 (167 bp, chr8: 141889570-141889736; hg19), which encodes a 55–amino acid peptide (G66–W120) forming part of the F1 lobe of the FERM domain (Fig.1A). We then quantified the PSI of the FAK lacking exon 4 (FAKΔe4) in normal and tumor tissues. Interestingly, FAKΔe4 mRNA was ubiquitously expressed across a wide range of tissues in healthy organs with a range of FAK exon 4 PSI between 0.913 and 0.472 (Fig.S2) (41). In addition, we showed that the FAK exon 4 PSI decreases in solid tumor tissues compared to the corresponding healthy tissues such as breast cancer (BRCA), colon adenocarcinoma (COAD), cholangiocarcinoma (CHOL), lung squamous cell carcinoma (LUSC), stomach adenocarcinoma (STAD), rectum adenocarcinoma (READ) and esophageal carcinoma (ESCA) using TCGA datasets (Fig.1B-H). Together, these data indicate that FAKΔe4 is overexpressed in numerous solid tumor types and may represent a clinically relevant factor.

**Fig. 1.**
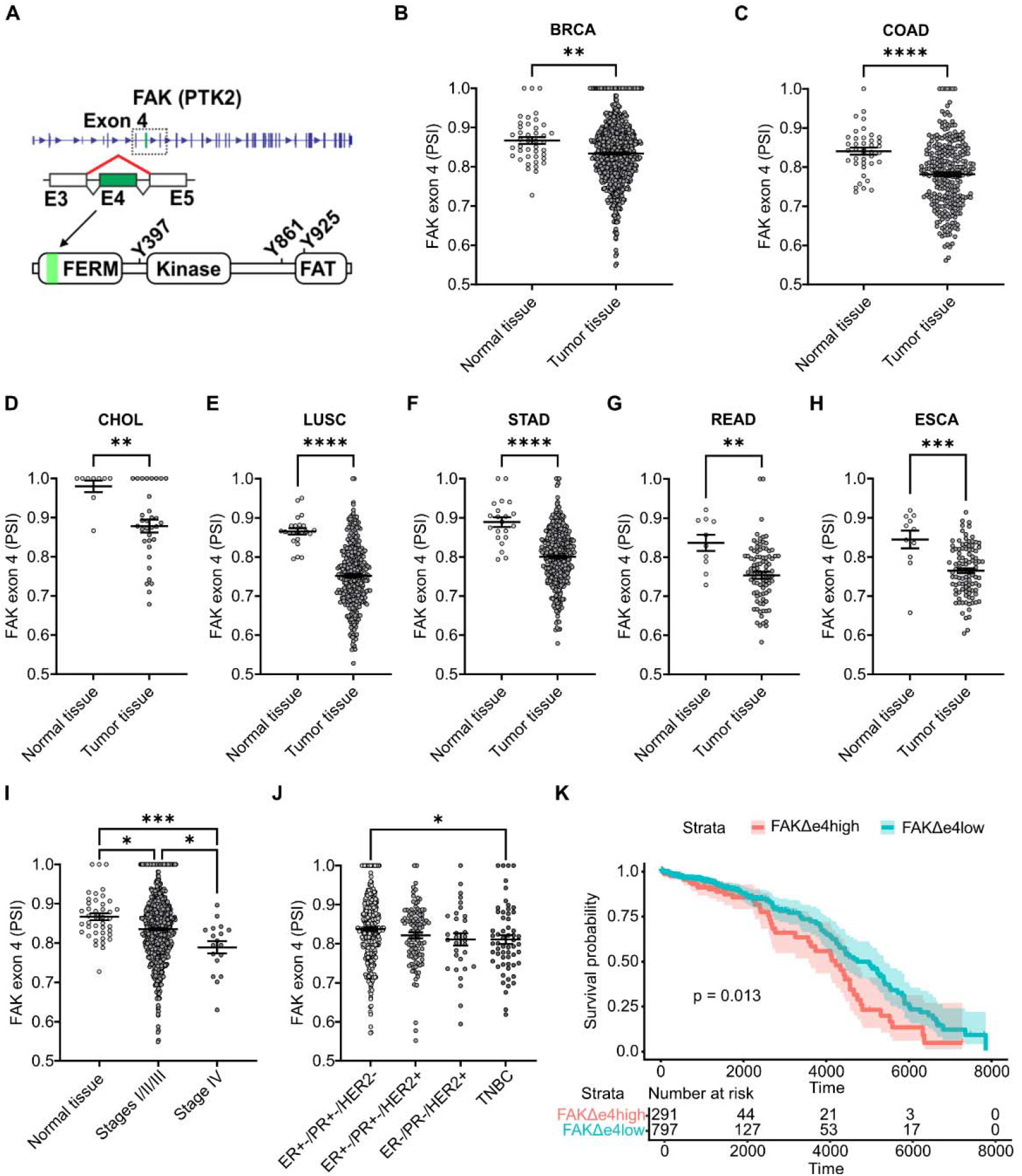
Detection of FAKΔe4 in different normal tissues and solid tumors. (**A**) Schematic view of FAK (PTK2) and the exon 4 splicing event that generates the FAKΔe4 protein. (**B**) Quantification of FAK alternative exon 4 splicing presented as percent spliced-in (PSI) from TCGA datasets of normal and cancerous tissues, including breast cancer (BRCA), (**C**) colon adenocarcinoma (COAD), (**D**) cholangiocarcinoma (CHOL), (**E**) lung squamous cell carcinoma (LUSC), (**F**) stomach adenocarcinoma (STAD), (**G**) rectum adenocarcinoma (READ) and (**H**) esophageal carcinoma (ESCA). Unpaired student *t* test. Data are means ± SEM; **p<0.01, ***p<0.001, ****p<0.0001. (**I**) Quantification of FAK alternative exon 4 percent spliced-in (PSI) in normal breast tissue, non-metastatic breast cancer stages (I/II/III) and metastatic breast cancer stage (IV) using BRCA-TCGA datasets. (**J**) Quantification of FAK alternative exon 4 PSI as a function of BRCA molecular subtypes from better to worst prognosis (ER+-/PR+- /HER2-, ER+-/PR+-/HER+, ER-/PR-/HER2+, TNBC). Data are means ± SEM; *p<0.05, ***p<0.001. Kruskal-Wallis test. (**K**) Corresponding Kaplan-Meier survival curves of BRCA patients grouped into lowFAKΔe4 expression (blue) and high FAKΔe4 expression (red).

To further explore the potential clinical relevance of this uncharacterized splice variant, we focused on breast cancer because of its established relationship between tumor progression and tumor stiffening processes (42,43). We found that FAKΔe4 expression increased progressively from normal tissue to non-metastatic tumors (stages I-III) and then to metastatic tumors (stage IV) (Fig.1I). We next analyzed FAKΔe4 expression across breast cancer molecular subtypes from better to worst prognosis: ER+-/PR+-/HER2+, ER+-/PR+-/HER2-, ER-/PR-/HER2+ and triple negative breast cancer (TNBC). We observed that FAKΔe4 is more expressed in the most aggressive group (TNBC) compared to the least aggressive group (ER+-/PR+-/HER2+) (Fig.1J). Finally, we assessed whether FAKΔe4 expression correlates with patient survival by stratifying patients into low or high expression groups. Importantly, patients with high FAKΔe4 expression showed reduced survival compared to those with low expression levels, indicating that elevated FAKΔe4 expression is associated with poorer clinical outcome (Fig.1K). Together, these analyses suggest that FAKΔe4 expression correlates with breast cancer progression, tumor aggressiveness, and worse prognosis.

### FAK**Δ**e4 is mechanoregulated

ECM stiffness has been reported as a regulator of alternative splicing (30,33). Thus, we investigated whether ECM stiffness regulates FAK alternative splicing. First, we explored if the FAK exon 4 PSI was regulated by tumor stiffness in TCGA breast cancer patient datasets. Since ECM stiffness measurements are not available in the public TCGA RNA-sequencing datasets, we used a 14-gene signature downstream of the YAP mechanosensing response as a proxy for tumor stiffness (30). Interestingly, the FAK exon 4 PSI followed a biphasic distribution according to the YAP signature score, with a minimal peak PSI around 0.8 in breast cancer patient datasets (Fig.2A). To confirm that ECM stiffness regulates FAKΔe4 splicing, we analyzed two public RNA sequencing datasets of malignant breast cell lines (MCF10CA1a and MDA-MB-231) cultured on different stiffnesses (GEO accession numbers: GSE205816 and GSE255829). The MCF10CA1a cells expressed more FAKΔe4 when cultured on stiff condition (8 kPa) compared to soft condition (0.8 kPa) (Fig.2B). Moreover, the metastatic breast cancer MDA-MB-231 cells also expressed more FAKΔe4 when cultured on stiff condition (308 kPa) compared to soft condition (1 kPa) (Fig.2C). We next performed long-read RNA sequencing on premalignant MCF10a and aggressive breast cancer MDA-MB-231 cell lines cultured on 1 and 10 kPa. In addition to confirming that FAKΔe4 encoded for a full-length transcript, the results suggested that FAKΔe4 is more expressed at 1 kPa in MCF10a compared to 10 kPa whereas FAKΔe4 is more expressed at 10 kPa in MDA-MB-231 compared to 1 kPa (Fig.S3A-D). In light of those results, we expanded the range of substrate stiffness (0.5 kPa to ∼70 GPa) to investigate and validate by RT-PCR whether FAKΔe4 splicing exhibited a biphasic response as a function of stiffness in MCF10a, malignant MCF7 and MDA-MB-231 cells. Similar to the biphasic distribution of FAKΔe4 splicing in patient samples, FAKΔe4 mRNA expression exhibited a biphasic relationship with substrate stiffness in MDA-MB-231, MCF7 and MCF10a cells with a maximum expression at 10 (1.36-fold increase), 5 (1.2-fold increase) and 1 kPa (1.31-fold increase), respectively (Fig.2D-E). Since alternative splicing is regulated by ECM stiffness through cell contractility (9,33), we investigated to which extent FAKΔe4 expression depends on actomyosin contractility. MDA-MB-231 cells were cultivated on 10 kPa to obtain the maximum FAKΔe4 expression and treated with the ROCK inhibitor Y27632 or myosin II inhibitor blebbistatin for 24 hours. We observed that both Y27632 and blebbistatin decreased FAKΔe4 expression compared to non-treated MDA-MB-231 cells (Fig.2F). We then sought to confirm whether the stiffness-mediated FAKΔe4 expression we observed could be detected at the protein level. Interestingly, a polyclonal anti-FAK antibody revealed two bands at 115 and 125 kDa for MDA-MB-231 cells cultured on 1 kPa and 10 kPa (Fig.2G). Quantification of the lower band signal (expected FAKΔe4 molecular weight) revealed a fold increase of 1.32 from 1 kPa to 10 kPa (Fig.2H), matching the expression increase observed at the mRNA level (Fig.2E). To demonstrate that the lower band correspond to the FAKΔe4 isoform, we implemented a strategy based on the use of a monoclonal anti-FAK antibody (FAK clone 4.47) whose target epitope is located in the N-terminus region of FAK. The ability of this antibody clone to detect only canonical FAK and not FAKΔe4 was confirmed by performing western blots on cells expressing an eGFP-tagged construct of the FAK isoforms (eGFP-cFAK or eGFP-FAKΔe4) (Fig.S4A). Next, considering the overlap of the 115 and 125 kDa bands, we performed a 2D electrophoresis to properly separate endogenous canonical FAK and endogenous FAKΔe4 of MDA-MB-231 cells cultured on 10 kPa again to ensure optimal FAKΔe4 expression. Two spots were visible at 125 kDa (pH≈6.2) and 115 kDa (pH≈6.5) after using the polyclonal anti-FAK (Fig.S4B). In contrast, an incubation with the FAK clone 4.47 antibody only revealed the spot at 125 kDa, which corresponds to the canonical FAK, while the spot at 115 kDa that would match FAKΔe4 was absent (Fig.S4B). Importantly, no additional FAK isoforms matching this molecular weight were detected in our RNA-seq analysis, strongly suggesting that the protein detected at 115 kDa is indeed FAKΔe4. Overall, these results indicate that FAKΔe4 is regulated by ECM stiffness in a biphasic manner, both *in vitro* and *in situ,* and is controlled through actomyosin contractility.

**Fig. 2.**
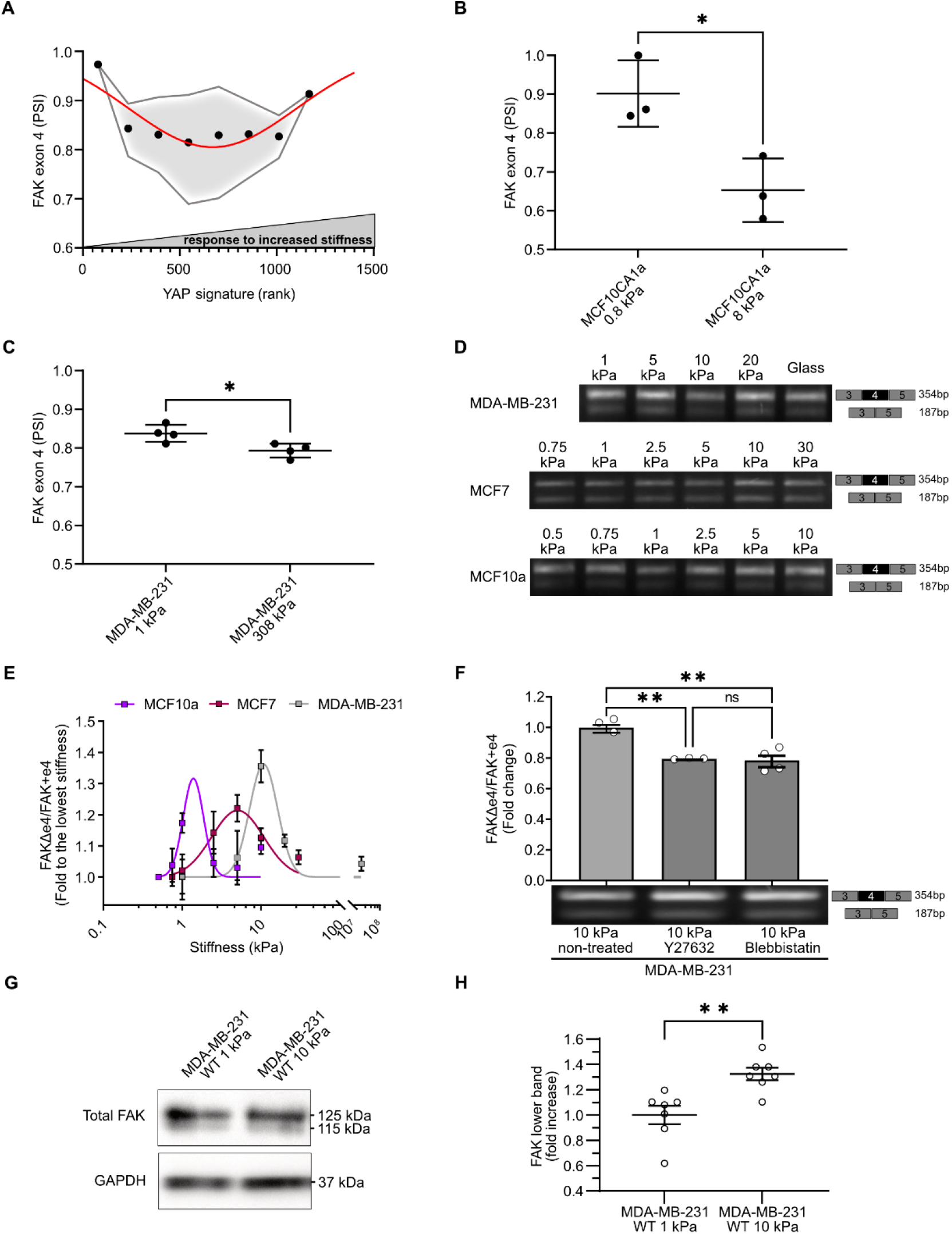
FAKΔe4 splice variant is mechanoregulated. (**A**) Quantification of FAK exon 4 percent spliced-in (PSI) in breast cancer patients (TCGA) as a function of a YAP signature used as a proxy of tumor stiffness. Data were fitted with a gaussian curve, and the envelope shows the adjusted variance of the PSI distribution. (**B**) Quantification of FAK exon 4 PSI in MCF10CA1a cells cultured on 0.8 and 8 kPa. Data analyzed from GSE205816. (**C**) Quantification of FAK exon 4 PSI in MDA-MB-231 cells cultured on 1 and 308 kPa. Data analyzed from GSE255829. (**D**) RT-PCR assays validating the stiffness-mediated splicing of FAK exon 4. The PCR products either include or exclude FAK exon 4 according to a range of stiffness: 1, 5, 10, 20 kPa and glass (≈70 GPa) in MDA-MB-231 cells, 0.75, 1, 2.5, 5, 10, 30 kPa for MCF7 cells and 0.5, 0.75, 1, 2.5, 5, 10 kPa for MCF10a cells. (**E**) Corresponding densitometric quantification of the FAKΔe4/FAK+e4 expression ratio determined by RT-PCR as function of substrate stiffness in MCF10a, MCF7 and MDA-MB-231 cells (N=6). Data normalized to the lowest stiffness. Data points were fitted with a gaussian curve to measure the optimal stiffness. (**F**) RT-PCR assay with the mRNA of MDA-MB-231 cells cultured on 10 kPa and treated with Y27632, blebbistatin or vehicle (non-treated, NT) showing FAK including or excluding exon 4. Corresponding densitometric quantification of the fold change of FAKΔe4/FAK+e4 ratio to the NT condition determined by RT-PCR (N=3). (**G**) Western blotting of whole protein extracts from MDA-MB-231 cells seeded on 1 and 10 kPa substrates showing the total FAK chemiluminescent signal and GAPDH as the loading control. (**H**) Corresponding densitometric quantification of FAK lower band expression for MDA-MB-231 cells on 1 kPa and 10 kPa as a fold increase baseline (N=7). Data normalized to the lowest stiffness. Data are means ± SEM; *p<0.05, **p<0.01.

### FAK**Δ**e4 regulates stiffness-driven cell spreading, migration and invasion

FAK is a key mechanosensing protein involved in regulating stiffness-driven cell behaviors such as spreading and migration (9,44,45). In this context, we investigated what was the contribution of the FAKΔe4 isoform in these processes. Since FAKΔe4 is always co-expressed with the canonical FAK including exon 4, we went for a strategy where we could measure the impact of different canonical FAK/FAKΔe4 ratio. We generated MDA-MB-231 cell populations with known FAKΔe4/FAK ratios by sorting cells expressing the eGFP-tagged canonical FAK or FAKΔe4 gated on the fluorescent intensity of each cell (Fig.S5A-B). MDA-MB-231 cells with high eGFP-cFAK expression were selected as the population where canonical FAK dominates with very low FAKΔe4 ratio (cFAK^high^), while MDA-MB-231 cells with mid-levels (FAKΔe4^mid^) and high-levels of eGFP-FAKΔe4 (FAKΔe4^high^) expression signal were selected to represent increased or very high FAKΔe4 ratio (Fig.S6A-B). We then investigated how the FAKΔe4 ratio influences the spreading area by seeding MDA-MB-231 cFAK^high^ or FAKΔe4^high^ on 0.5, 1, 5, 10, 18 and 50 kPa and measuring cell area after 24 hours. Our data revealed that the maximum cell spreading was reached on a substrate stiffness of 10 kPa for cFAK^high^ cells compared to 5 kPa for FAKΔe4^high^ cells (Fig.3A-B). Migration speed as a function of substrate stiffness is known to be biphasic in parental MDA-MB-231, with a maximum speed achieved between 10 and 18 kPa (11). We assessed the cell migration speed in MDA-MB-231 cFAK^high^ or FAKΔe4^high^ cells as a function of stiffness (Fig.3C). We found that the optimal stiffness for cell speed was between 10 and 18 kPa for cFAK^high^ cells compared to 5 kPa for FAKΔe4^high^ cells (Fig.3D). Moreover, after reaching its maximum, the migration speed of the FAKΔe4^high^ cells dropped rapidly to levels below the speed observed for cFAK^high^ cells, even for the stiffest condition used. We also quantified the mean square displacement (MSD) as a quantitative measurement of cell displacement and scattering during migration over time (46). Surprisingly, we observed that cFAK^high^ cells have a greater directional persistence on 1 and 5 kPa, whereas FAKΔe4^high^ cells have a greater directional persistence on 10 and 18 kPa (Fig.3E). Next, we expanded our analysis to 3D cell migration and invasion by embedding cFAK^high^ or FAKΔe4^high^ spheroids into 1.5 mg/mL collagen gels that were untreated (≈180 Pa) or pre-crosslinked using non-enzymatic glycation with 100 mM ribose (≈500 Pa) (47). Cell invasion was then quantified over 3 days. Interestingly, FAKΔe4^high^ cells displayed increased invasion ability in the softer collagen gel compared to cFAK^high^ cells (Fig.3F-H). However, this difference is lost in the stiffer collagen gel as the measured invasion of cFAK^high^ cells increased while decreased in FAKΔe4^high^ cells to reach similar levels (Fig.3G-H). Finally, we tested the impact on collective cell migration by performing wound healing assays of cFAK^high^ and FAKΔe4^high^ cells cultured on collagen-coated 96-well plates. In this case, the wound closure was slightly slower in FAKΔe4^high^ cells compared to cFAK^high^ cells (Fig.S7A-B), which is consistent with our migration result on the higher stiffness. Taken together, FAKΔe4 appears to upregulate cell spreading and migration at lower substrate stiffness in both 2D and 3D contexts.

**Fig. 3.**
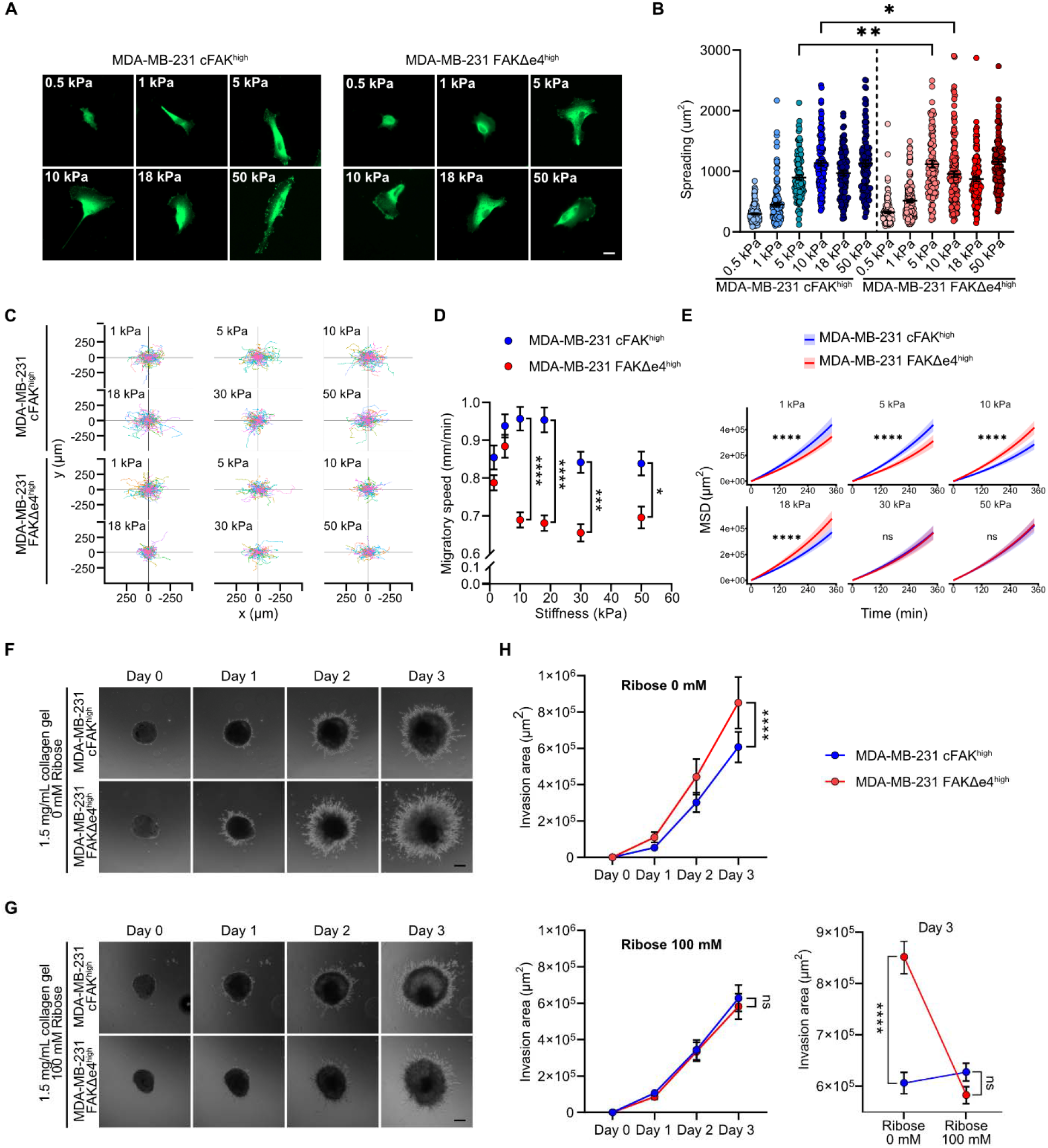
FAKΔe4 regulates cell spreading, migration and invasion in a stiffness-dependent manner. (**A**) Representative immunofluorescence images of MDA-MB-231 cFAK^high^ and FAKΔe4^high^ cells cultured on 0.5, 1, 5, 10, 18 and 50 kPa using confocal microscopy. Scale bar = 10 μm. (**B**) Corresponding fluorescence quantification showing cell spreading of MDA-MB-231 cFAK^high^ and FAKΔe4^high^ cells cultured on 0.5, 1, 5, 10, 18 and 50 kPa. N=3. (**C**) Tracks from individual MDA-MB-231 cFAK^high^ and FAKΔe4^high^ cells cultured on 1, 5, 10, 18, 30 and 50 kPa. N=3. (**D**) Corresponding quantification of cell migration speed of MDA-MB-231 cFAK^high^ and FAKΔe4^high^ cells cultured on 1, 5, 10, 18, 30 and 50 kPa. (**E**) Modeling of the mean square displacement (MSD) according to time comparing both MDA-MB-231 cFAK^high^ and FAKΔe4^high^ cells on each stiffness (1, 5, 10, 18, 30 and 50 kPa). (**F**) Representative images of spheroid outgrowth of MDA-MB-231 cFAK^high^ and FAKΔe4^high^ cells at 24, 48 and 72 hours post-embedding in 1.5 mg/mL 3D collagen gels with 0 mM ribose. Scale bar = 100 μm. (**G**) Representative images of spheroid outgrowth of MDA-MB-231 cFAK^high^ and FAKΔe4^high^ cells at 24, 48 and 72 hours post-embedding in 1.5 mg/mL 3D collagen gels with 100 mM ribose. Scale bar = 100 μm. (**H**) Corresponding quantification of invaded area of MDA-MB-231 cFAK^high^ and FAKΔe4^high^ spheroids with 0 mM and 100 mM ribose after 24, 48 and 72 hours. N=5. Two-way ANOVA with Tukey. Data are means ± SEM; ns=non-significant, *p<0.05, **p<0.01, ***p<0.001, ****p<0.0001.

### FAK**Δ**e4 impairs focal adhesion dynamics

Cell migration and invasion depend on a tight regulation of FA dynamics and turnover (48,49). In fact, FAK residency time within FA has been associated with FA turnover (50). Therefore, we performed a Fluorescence Recovery After Photobleaching (FRAP) experiment to measure whether the residency time of FAKΔe4 within FAs was different from the canonical FAK. Single FAs were photobleached and the fluorescence was followed in real time for MDA-MB-231 cFAK^high^ and FAKΔe4^high^ cells (Fig.4A). The recovery of eGFP-FAKΔe4 at FAs was altered compared to eGFP-cFAK (Fig.4A, B, C). In addition, we measured the plateau reached as for the final recovery level to assess the mobile and immobile fractions of FAK. About 50% of eGFP-FAKΔe4 pool within the FAs was mobile while the eGFP-cFAK mobile fraction in the FAs was 80% (Fig.4C-D). We then investigated whether FAKΔe4 impacted FA turnover by performing an 80-min live imaging assay using total internal reflection fluorescence (TIRF) microscopy (Fig.4E). At the single FA level, we assessed FA assembly and disassembly rate, stability and lifetime. FAKΔe4-containing FAs displayed a lower FA assembly and disassembly rate compared to cFAK-containing FAs (Fig.4F-G). Moreover, FAKΔe4-containing FAs were more stable and had longer lifetime compared to cFAK-containing FAs (Fig.4H-I). Of note, both FAKΔe4 and cFAK were strongly colocalized with total FAK and highly enriched in FAs (Fig.S8A-B), indicating that FAKΔe4 intracellular localization remained unchanged. Together, these data show that FAKΔe4 strongly influences FA dynamics and turnover.

**Fig. 4.**
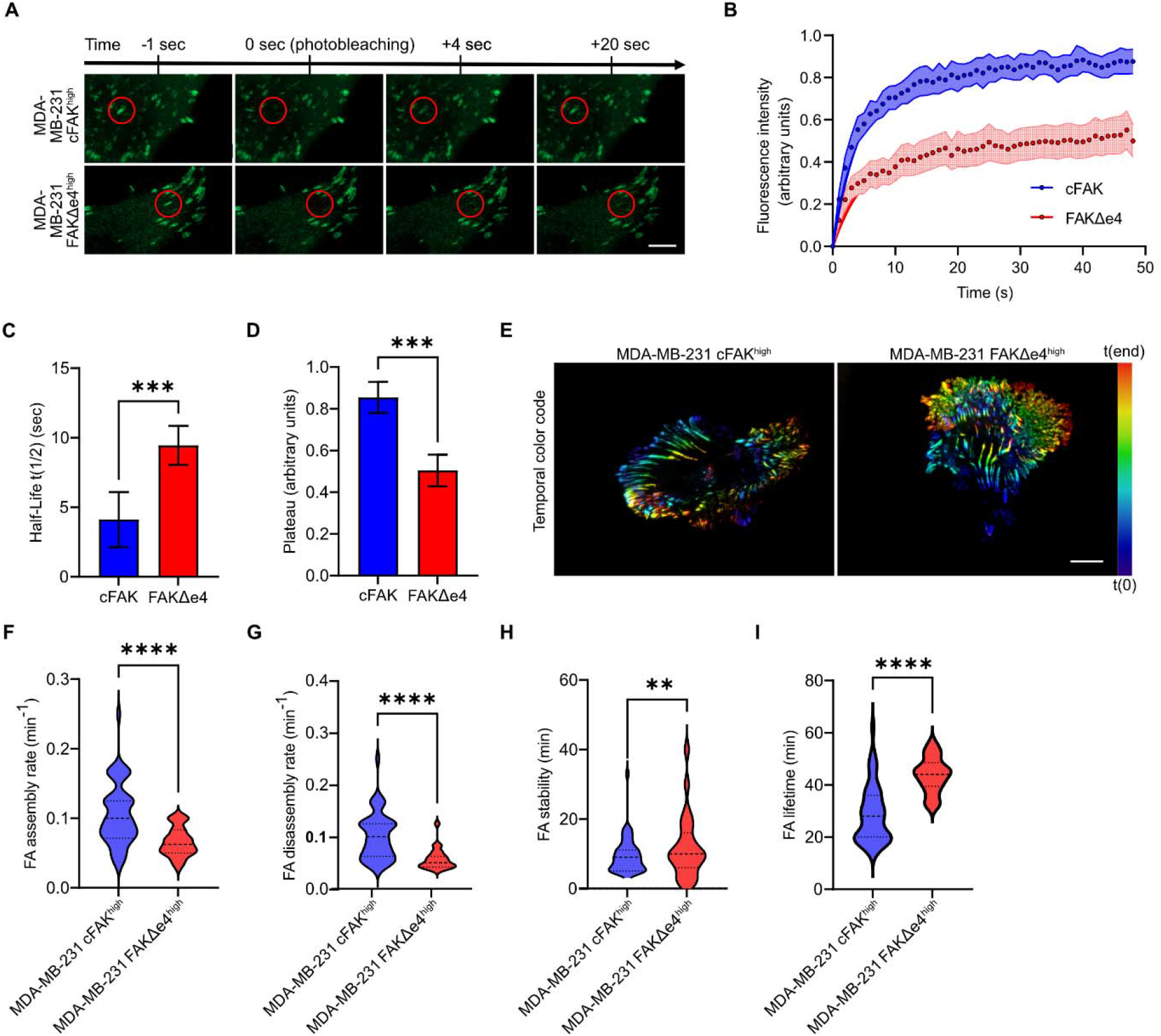
FAKΔe4 impacts focal adhesion dynamics. (**A**) Representative images from timelapse during the Fluorescence Recovery After Photobleaching (FRAP) experiment showing different steps: before photobleaching, photobleaching at a single FA, and the fluorescence recovery after 4 and 20 sec of MDA-MB-231 cFAK^high^ and FAKΔe4^high^ cells cultured on glass. The red circles show the targeted FA in MDA-MB-231 cFAK^high^ and FAKΔe4^high^ cells. Scale bar = 5 μm. (**B**) Quantification of eGFP-cFAK and eGFP-FAKΔe4 fluorescence recovery following FA photobleaching in MDA-MB-231 cells. The normalized mean intensity is shown along with the exponential fit. (**C**) Corresponding quantifications of the recovery half-life and (**D**) of the immobile fraction plateau measured from the FRAP analysis of eGFP-cFAK and eGFP-FAKΔe4 cells. N=3. Student t-test. Data are means ± SEM; ***p<0.001. (E) Temporal color-coded images generated from an 80-min timelapse of FAs of MDA-MB-231 cFAK^high^ and FAKΔe4^high^ cells cultured on glass (1 image per 120 seconds) using TIRF microscopy. Scale bar = 10 μm. (F) Corresponding quantifications of the FA assembly rate, (**G**) FA disassembly rate (**H**) FA stability and (**I**) FA lifetime. Student t-test. Data are median ± quartile; **p<0.01, ***p<0.001, ****p<0.0001.

### FAK**Δ**e4 decreases focal adhesion maturation state

FA number, size and maturation collectively influence cell migration (21,22,51). Given our results showing that FA dynamics and turnover were impacted by FAKΔe4, we sought to determine if FA maturation was also altered. Since the gamut of FA size reflects the continuum of FA maturation (52,53), we first quantified the number and size distribution of FAs in MDA-MB-231 cFAK^high^ and FAKΔe4^high^ cells. We observed that FAKΔe4^high^ cells had a higher number of FAs per cell compared to cFAK^high^ cells (Fig.5A-B). In addition, a larger proportion of FAs were smaller in FAKΔe4^high^ cells compared to those in cFAK^high^ cells (Fig.5C). These data suggest that the FA maturation process could be altered in the presence of FAKΔe4. FA maturation involves the sequential recruitment of structural proteins such as talin, vinculin and zyxin (54,55). Talin is recruited in nascent adhesions before FAK recruitment, while vinculin is recruited following FAK recruitment during the mid-term stage of FA maturation, and zyxin during the late stage of FA maturation (54–56). We investigated by immunofluorescence whether FAKΔe4 altered the recruitment of those proteins to FAs. First, we showed that talin colocalizes similarly with cFAK and FAKΔe4 (Fig.5D-E). On the other hand, vinculin and zyxin colocalization with FAKΔe4 is decreased compared to cFAK (Fig.5F-I). Together, these results suggest that FAKΔe4 leads to the formation of increased number but smaller FAs, which is consistent with the impairment of FA maturation process.

**Fig. 5.**
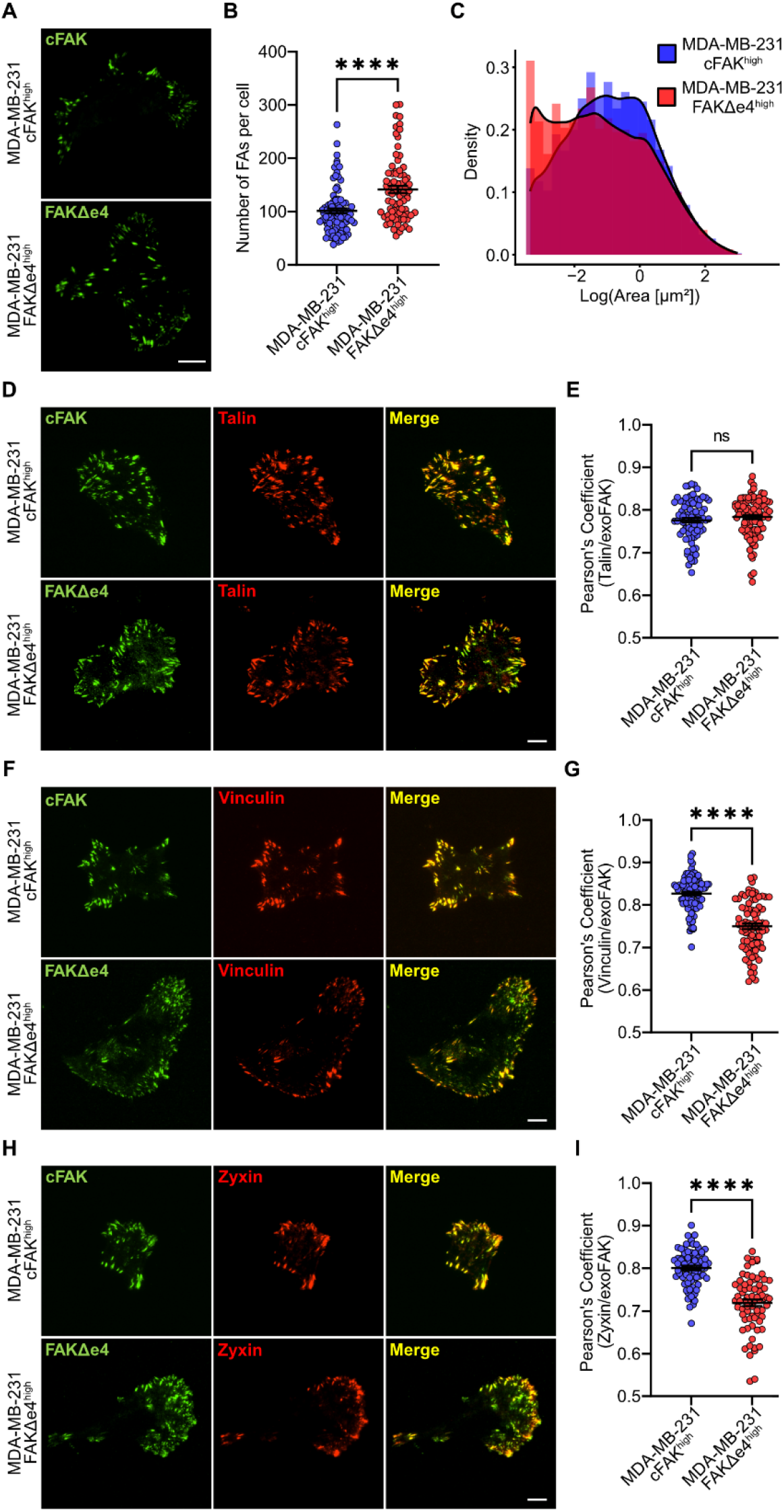
FAKΔe4 influences focal adhesion maturation. (**A**) Representative images of MDA-MB-231 cFAK^high^ and FAKΔe4^high^ cells cultured on glass using TIRF microscopy. (**B**) Corresponding quantification of the number of FAs per cell comparing MDA-MB-231 cFAK^high^ and FAKΔe4^high^ cells. N=3. (**C**) Corresponding density quantification of the FA number according to their size comparing MDA-MB-231 cFAK^high^ and FAKΔe4^high^ cells. Wilcox test. (**D**) Representative TIRF microscopy images of anti-eGFP (exogenous cFAK or FAKΔe4) and anti-talin stainings in MDA-MB-231 cFAK^high^ and FAKΔe4^high^ cells. (**E**) Corresponding quantification of colocalization using Pearson’s Coefficient in MDA-MB-231 cFAK^high^ and FAKΔe4^high^ cells comparing talin vs. exogenous FAK (cFAK or FAKΔe4). N=3. (**F**) Representative TIRF microscopy images of anti-eGFP (exogenous cFAK or FAKΔe) and anti-vinculin stainings in MDA-MB-231 cFAK^high^ and FAKΔe4^high^ cells. (**G**) Corresponding quantification of colocalization using Pearson’s Coefficient in MDA-MB-231 cFAK^high^ and FAKΔe4^high^ cells comparing vinculin vs. exogenous FAK (cFAK or FAKΔe4). N=3. (**H**) Representative TIRF microscopy images with the anti-eGFP (exogenous cFAK or FAKΔe) and anti-zyxin stainings in MDA-MB-231 cFAK^high^ and FAKΔe4^high^ cells. (**I**) Corresponding quantification of colocalization using Pearson’s Coefficient in MDA-MB-231 cFAK^high^ and FAKΔe4^high^ cells comparing zyxin vs. exogenous FAK (cFAK or FAKΔe4). N=3. Student t-tests. Data are means ± SEM; ns=non-significant, ****p<0.0001. Scale bars = 10 μm.

### FAKΔe4 drives cell mechanotransduction responses at cellular and molecular levels

FAK Y397 autophosphorylation is a key mechanotransduction event in cells (9). In this context and given our results on migration and FAs in FAKΔe4^high^ cells, we asked if FAKΔe4 mechanotransduction response was altered compared to cFAK. First, MDA-MB-231 cFAK^high^ and FAKΔe4^high^ cells were cultured on 1 and 10 kPa for 48 hours before assessing pY397-FAK by Western blot (Fig.6A). Quantification of pY397-FAK normalized over the total FAK revealed that FAKΔe4 was more phosphorylated on 1 kPa compared to cFAK (Fig.6A-B). However, normalized FAKΔe4 and cFAK pY397 levels were found to be equivalent in cells seeded on 10 kPa (Fig.6A-C). To further confirm this result, we expanded our investigation to stiffness-mediated YAP nuclear translocation for cells seeded on substrate of 0.5 to 50 kPa (Fig.6D). Interestingly, while both cFAK^high^ and FAKΔe4^high^ cells displayed increased YAP nuclear translocation as a function of stiffness up to 5 kPa, YAP nuclear translocation continued to increase in cFAK^high^ until 10 kPa before decreasing at 18 kPa compared to FAKΔe4^high^ in which YAP nuclear signal peaked at 5 kPa and then decreased at 10 and 18 kPa (Fig.6E). In both cases, the YAP nuclear signal appeared to increase again for stiffness above 18 kPa (Fig.6E). Furthermore, FAK activation is required for cell durotaxis in MDA-MB-231 cells (57). Therefore, we investigated if FAKΔe4 influences durotactic behavior by measuring cell density distribution 3 days after seeding the cells on hydrogels presenting stiffness gradient ranging from 1 to 30 kPa as described before (17). Since these are continuous gradients with very high spatial stiffness resolution that can resolve small difference in durotactic behavior (17), we proceeded to test the parental MDA-MB-231, cFAK^high^, FAKΔe4^mid^ and FAKΔe4^high^ cells. First, the cell density distribution of parental MDA-MB-231 cells peaked at an optimal stiffness of 16.3 kPa (Fig.S9), providing a baseline for the durotactic behavior of these cells consistent with their known optimal cell migration speed. The optimal peak cell density in cFAK^high^ cells was found to occur at a stiffness of 18.2 kPa (Fig.6F, I), which was close to the value of the parental population. In contrast, FAKΔe4^mid^ and FAKΔe4^high^ cells optimal peak migration were measured at 14.4 kPa and 8.5 kPa, respectively (Fig.6G-I). Taken together, our results identify FAKΔe4 as a molecular switch that facilitates mechanotransduction response at lower ECM stiffness, which in turn influences how cells sense and find an optimal substrate stiffness, notably during durotaxis.

**Fig. 6.**
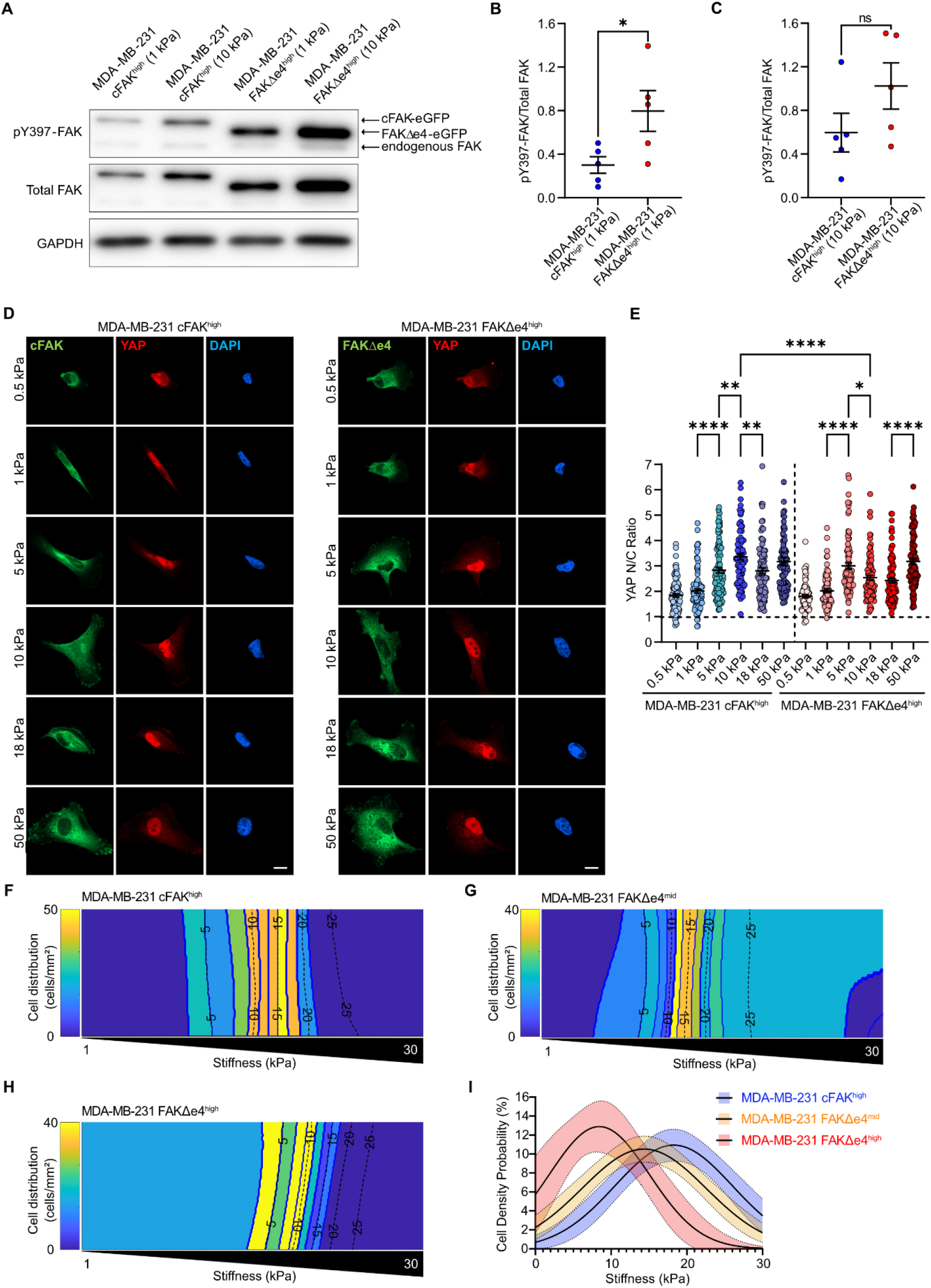
FAKΔe4 drives cell mechanotransduction responses at cellular and molecular levels. (**A**) Western blots of whole protein extracts from MDA-MB-231 cFAK^high^ and FAKΔe4^high^ cells seeded on 1 and 10 kPa substrates. GAPDH was used as loading control. (**B**) Corresponding quantification of pY397-FAK/total FAK ratio for MDA-MB-231 cFAK^high^ and FAKΔe4^high^ cells seeded on 1 kPa. N=5. (**C**) Corresponding densitometric quantification of pY397-FAK/total FAK ratio for MDA-MB-231 cFAK^high^ and FAKΔe4^high^ cells seeded on 10 kPa. N=5. Student t-tests. (**D**) Representative immunofluorescence images showing MDA-MB-231 cFAK^high^ and FAKΔe4^high^ cells cultured on 0.5, 1, 5, 10, 18 and 50 kPa stained for eGFP (exogenous FAK) in green, YAP in red and DAPI in blue. Scale bars = 10 μm. (**E**) Corresponding quantification of YAP nuclear/cytoplasm ratio in MDA-MB-231 cFAK^high^ and FAKΔe4^high^ cells cultured on 0.5, 1, 5, 10, 18 and 50 kPa. The exogenous FAK channel was used to determine the cytoplasm edge and the DAPI channel was used to determine the nucleus edge (N=3). One-way ANOVA with Tukey. (**F**) Representative heat maps of cell density probability as a function of substrate stiffness between 1 to 30 kPa for MDA-MB-231 cFAK^high^, (**G**) FAKΔe4^mid^ and (**H**) FAKΔe4^high^ cells. (**I**) Corresponding quantifications of the cell density probability according to substrate stiffness presented as a Gaussian fit with 95% confidence interval for cFAK^high^ (N=6), FAKΔe4^high^ (N=3) and FAKΔe4^mid^ (N=3) cells. Extra sum-of-squares F-test on gaussian curve means. Data are means ± SEM; ns=non-significant, *p<0.05, **p<0.01 and ****p<0.0001.

## Discussion

While ECM stiffness has been shown to alter alternative splicing events through cell mechanotransduction, the role of stiffness-mediated alternative splicing on cell mechanosensing remains unknown. Here, we demonstrated the existence of an interplay between ECM stiffness, alternative splicing and mechanosensing. We showed that FAKΔe4 is expressed ubiquitously across healthy tissues while being overexpressed in numerous solid tumors. More specifically, FAKΔe4 is overexpressed according to tumor stages and molecular subtype classification in breast cancer in addition to being a potential clinically relevant factor. We determined that FAKΔe4 is regulated in a biphasic manner by substrate stiffness and cell contractility. Importantly, we demonstrated that FAKΔe4 lowers the stiffness at which optimal cell spreading, migration and invasion occur. Furthermore, we showed that FAKΔe4 alters FA dynamics and prevent their maturation. Critically, FAKΔe4 displayed facilitated stiffness-mediated activation which was also associated with a shift in the ECM stiffness required for YAP nuclear translocation and durotaxis homing. Together, we revealed that FAKΔe4 is regulated by ECM stiffness and, in turn, serves as a mechanosensor switch for tumor cells to fine-tune cell mechanosensing.

FAK is a mechanosensitive protein which depends on force-induced conformational change to be activated through autophosphorylation (9,20). Mechanistically, in closed conformation, the FERM domain of FAK binds the kinase domain acting as an autoinhibitory domain preventing Y397 from autophosphorylation (58). Strikingly, the FAK exon 4 encodes for a 55–amino acid peptide (G66–W120) forming part of the F1 lobe of the FERM domain. The absence of FAK exon 4 peptide may play a critical role in FAK autophosphorylation as D101 and H99 interact with the Y397-containing FAK linker situated between the FERM domain and kinase domain (59). The absence of FAK exon 4 could weaken or disrupt the autoinhibitory interactions between the FERM, kinase and linker domains, facilitating FAK activation. This would explain the overactivation of FAKΔe4 in cells cultured onto low stiffness substrate presenting itself as constitutively active or in need of less mechanical force to be activated. Interestingly, FAK mutants with different N-terminal deletions promote FAK phosphorylation (60–62). In fact, the deletion of the first 124 amino acids was sufficient to result in increased FAK phosphorylation, something strikingly similar to our present observation with FAKΔe4 (62). In this context, FAK splicing can be viewed within a broader framework in which alternative splicing remodels protein activity and function.

FA maturation state is closely linked to FAK dynamics. Concurrently, longer time residency of FAK at FAs is associated to less mature FAs (63). Moreover, increased FAK Y397 phosphorylation promotes the time residency of FAK at FAs (23), which is in line with our current results. In addition, we observed that FAKΔe4 impairs FAs as shown by vinculin and zyxin recruitment. Interestingly, FAK Y397 phosphorylation is linked to the decision to proceed with FA maturation or disassembly (23). In both cases, paxillin appears to play a critical role. For instance, FAK mediates vinculin recruitment to FAs through paxillin phosphorylation (64). In fact, paxillin can act as an adaptor protein that links FAK to vinculin through FAK Y925 phosphorylation (65). This raise the possibility that FAKΔe4-derived impairment of FA maturation could be due to altered FAK-paxillin interaction. Alternatively, FAK regulates intracellular actomyosin forces through the activation of RhoA (66). Through these intracellular forces, vinculin is recruited to FAs by talin mechanical stretching which uncovers cryptic vinculin-binding domain (VBD) sites (67). Whether FAKΔe4 influences on FA dynamics occurs through altered downstream signaling (e.g. recruitment of signaling proteins such as paxillin), by altering the mechanical load on single FA sites or both, remains to be assessed. Nevertheless, the expression of FAKΔe4 acts as a modulator of FA dynamics and FA-related cellular functions such as migration.

In solid tumors, the ECM forms a negative stiffness gradient away from the edge of the tumor (68). Durotaxis has been defined as the directed migration of cells toward stiffer substrates. However, recent studies have redefined this concept, demonstrating that cells can migrate either toward softer or stiffer substrates in pursuit of an optimal stiffness (13,69). So far, some molecular regulators of durotaxis have been identified, but most of those identified were linked to FA regulation (13,70–73). This observation is consistent with a widespread model in mechanobiology known as the motor-clutch model, utilized to predict how cells respond to mechanical stiffness (74). The motor-clutch model predicts that debilitated FA maturation on stiff substrates will drive cells towards an optimal softer rigidity (75,76). This is coherent with cells expressing more FAKΔe4 displaying an optimal durotaxis stiffness that is lower than the canonical FAK. Tumor invasion into softer surrounding tissues requires that a subset of tumor cells must exhibit negative durotaxis (69). Thus, FAKΔe4 expression observed in tumors might serve as an adaptation that allows tumor cells to trigger this negative durotaxis once the ECM negative stiffness gradient reaches a certain stiffness level.

Contrary to other characterized FAK isoforms, we found that FAKΔe4 is expressed ubiquitously across both normal and tumor human tissues. In this context, FAKΔe4 could be involved in biological processes featuring transient ECM stiffening where FAK plays a critical role, including development, cell fate and wound healing (1,2,5,77). Moreover, the inducible loss of FAK in neural crest cells leads to craniofacial and cardiovascular developmental defects (78). Interestingly, the head mesoderm stiffens and affects the migration of neural crest cells during morphogenesis (79). As such, it is plausible that stiffness-mediated splicing of FAKΔe4 could play a role in triggering the cell migration in the neural crest. Moreover, FAK is a regulator of the wound healing efficiency in a stiffness-modulated environment (80). Intermediate optimal stiffness is required for cells to perform an effective wound healing mechanism, where a matrix that is either too soft or too stiff impedes the process (81). In this case, FAKΔe4 could serve as a mechanosensor to guide cell migration for wound closure in stiffness-dependent manner. Moreover, considering how FAKΔe4 optimal expression is cell-type specific as a function of stiffness, its expression could be linked to the physiological stiffness range of a tissue. Overall, FAKΔe4 could be an important driver of stiffness-dependent physiological processes.

Taken together, we demonstrated a mechanism based on the regulation of a FAK splice variant that allows cells to home toward their preferred stiffness. Importantly, the ubiquitous expression of FAKΔe4 across healthy tissues and tumor types highlights the major regulating role this splice variant could play in influencing migration during development, regeneration or cancer progression. These findings raise critical questions about our understanding of mechanotransduction and the importance of biphasic splicing events to fine-tune biological processes.

## Limitations of the study

One of the limitations of our study is the reliance on an overexpression strategy to determine how the FAK splice variant impacts cell behavior. While this strategy broadly represents the fact that FAKΔe4 co-exists endogenously with canonical FAK, it prevents us from exploring in detail how it impacts the downstream signaling events as we cannot distinguish its contribution over that of canonical FAK. Even assessing changes in potential binding partners could be challenging as FAK dimerizes in FAs meaning that immunoprecipitation assays to pulldown FAKΔe4 could likely also pulldown canonical FAK if they co-dimerize (82). Establishing FAK knockout cell lines combined with expressing FAK constructs would facilitate the investigation of the specific molecular mechanisms associated with the FAKΔe4 isoform. Such a strategy will be essential for future work aimed at determining the underlying FAKΔe4-specific downstream signaling that alters the FA maturation process.

FAKΔe4 is ubiquitous and its optimal expression was shown to be cell-type specific as a function of stiffness. However, TCGA RNA sequencing datasets come from bulk biopsies that involve many cell types as well as tumor cell subpopulations, and each one might have a specific stiffness-mediated FAKΔe4 expression response that adds noise and limits the precision of our comparison with the YAP-derived signature. In addition, evaluating the impact of FAKΔe4 on the optimal stiffness for cell responses was inherently limited by the number of variables and experimental conditions in play. Regarding experiments involving 2D polyacrylamide gels for instance, the choice of substrate stiffness had to cover the stiffness range observed in breast cancer. This greatly limits achievable statistical power in most experiments, which is why we had to limit ourselves to the cFAK^high^ and FAKΔe4^high^ cell populations in several instances. With respect to our 3D experiments, the range of stiffness achievable using the non-enzymatic ribose crosslinking method without altering the collagen hydrogel architecture remains limited (83). Further increasing the ribose concentration or altering collagen concentration would also modify the collagen architecture, which would also impact the 3D cell migration (47,83).

At this point, it is also unclear how FAKΔe4 activation is facilitated. Notably, FAK activation and signaling is dependent of force-mediated conformation changes (20). Thus, high resolution single protein stretching assays could inform on the mechanical thresholds required to trigger FAKΔe4 conformational opening under tensional force. In parallel, it would be instructive to precisely assess the mechanical load on single FA in relationship with FAK activation in the presence of FAKΔe4. While such experiments are extremely challenging to perform and require specialized equipment and methods, this knowledge could eventually help explain and predict cell-type specific stiffness responses.

Finally, we have yet to identify the upstream mechanism that regulates FAK exon 4 splicing. Future experiments will need to uncover which upstream signaling events and splicing factors are involved. Uncovering those could prove to be critical to tune FAKΔe4 expression, but also to understand how other biphasic biological events are regulated.

## Resource availability

### Lead contact

Further information and requests for resources and reagents should be directed to, and will be fulfilled by the lead contact, Francois Bordeleau

### Materials availability

Plasmid generated in this study are available from the lead contact with a completed material transfer agreement.

### Data and code availability

This paper analyzes existing, publicly available data, accessible from The Cancer Genome Atlas Program portal (https://portal.gdc.cancer.gov/) or the Gene Expression Omnibus (GEO) under accession numbers GSE334884, GSE205816 and GSE255829.

## Acknowledgments

This work was supported in part by a Canadian Institutes of Health Research (CIHR) Project grant (PLL-179768) to FB and a Canada’s New Frontiers in Research Fund – Exploration grant (NFRFE-2023-00674) to FB and SL. The CHU de Québec-Université Laval Research Center has been awarded a grant from the Fonds de recherche du Québec (FRQ; https://doi.org/10.69777/30641). MM and OC are recipients of FRQ Doctoral Scholarships (https://doi.org/10.69777/320189, https://doi.org/10.69777/2001362), Excellence Graduate Scholarships from the Université Laval Cancer Research Center and Desjardins Excellence Graduate Scholarships from the Fondation du CHU de Québec. JG is the recipient of a joint CIHR – Cancer Research Society Doctoral Award. OC was also a recipient of Doctoral Training Awards from the Université Laval LOEX Center and the Eye Disease Foundation. SL was a Junior 2 Research Scholar of the FRQ (https://doi.org/10.69777/296806). FB is a tier 2 Canada Research Chair in Tumor Mechanobiology and Cellular Mechanoregulation.

## Author contributions

Conceptualization: MM, DS, FB; Methodology: MM, JG, DS, FB; Formal analysis and investigation: MM, JG, OC, ET, WZV, EP, GK, FB; Visualization: MM, JG, OC, FB; Software: FB; Writing - original draft preparation: MM, FB; Writing - review and editing: MM, JG, OC, ET, WZV, EP, GK, SL, SH, DS, FB; Funding acquisition: SL, FB; Supervision: SL, SH, DS, FB.

## Materials and methods

### Reagents and antibodies

Primary antibodies used were as follows: mouse anti-GFP (#11-814-460-001, Roche), rabbit anti-FAK (#3285S, Cell Signaling), rabbit anti-phospho Y397 FAK (#3283S, Cell Signaling), FAK clone 4.47 (#05-537, MilliporeSigma), rabbit anti-talin (#14168-1-AP, Proteintech), rabbit anti-vinculin (#26520-1-AP, Proteintech), rabbit anti-zyxin (#Z4751, Sigma), rabbit anti-YAP1 (#14074, Cell Signaling) and rabbit anti-GAPDH (#5174, Cell Signaling). Secondary antibodies used were as follows: Alexa Fluor 488-conjugated to goat anti-rabbit (#A11034, Invitrogen), Alexa Fluor 488-conjugated to goat anti-mouse (#A11029, Invitrogen), Alexa Fluor 594-conjugated to goat anti-rabbit (#A11037, Invitrogen) and HRP-link goat anti-rabbit (#7074S, Cell Signaling).

### Cell culture

Highly metastatic MDA-MB-231 breast adenocarcinoma cells, all modified MDA-MB-231 (#HTB-26, ATCC) cell lines and MCF7 breast cancer cells (#HTB-22, ATCC) were maintained in DMEM (#319-015-CL, Wisent Bioproducts) supplemented with 10% fetal bovine serum (FBS) (#A5256701, Thermo Fisher Scientific) and 1% penicillin-streptomycin (#450-201-EL, Wisent Bioproducts). MCF10a mammary epithelial cells (#CRL-10317, ATCC) were maintained in DMEM/F12 (#319-085-CL, Wisent Bioproducts) supplemented with 5% horse serum (#16050130, Thermo Scientific), 20 ng.mL^-1^ epidermal growth factor (EGF) (#5018597, Invitrogen), 10 mg.mL^-1^ insulin (#I6634, Sigma-Aldrich), 0.5 mg.mL^-1^ hydrocortisone (#HO888, Sigma-Aldrich), 100 ng.mL^-1^ cholera toxin (#C8052, MilliporeSigma) and 1% penicillin–streptomycin (#450-201-EL, Wisent Bioproducts). All cells were cultured at 37 °C and 5% CO_2_. For pharmacological inhibition of cell contractility, MDA-MB-231 cells were allowed to grow 48 hours on 10 kPa substrate following the addition of Y27632 (final concentration: 10μM) (#129830-38-2, Cayman Chemical Company) or blebbistatin (10μM) (#20339, Calbiochem) for 24 hours before cell lysis. For the non-treated condition (NT), an equivalent amount of DMSO to the vehicle was added.

### Reverse transcription PCR

MDA-MB-231, MCF7 and MCF10a cell lines were cultured on polyacrylamide (PA) gels for 72 hours, 72 hours or 48 hours, respectively. Samples were rinsed with PBS and collected after being treated with 0.05% trypsin solution. Then, RNA was extracted from cells using Total RNA Isolation Miniprep Kit EZ-10 Spin Column (#BS1361, Bio Basic). The reverse transcriptase step was performed using MultiScribe™ Reverse Transcriptase kit (#43-112-35, Thermo Fisher Scientific). PCR was performed using Taq DNA Polymerase with 10 mM dNTP Mix (#E00101, GenScript). A 3% agarose gel (#A436-07, J.T Baker) was used to resolve PCR products. The PCR primer set used was: forward primer (GCCAACCACCTGGGCCAGTATTAT) and reverse primer (TCCTGGTCCACTTGATCAGCTATCTCTAAC) to amplify both cDNA containing FAK exon 4 and cDNA excluding FAK exon 4 in the same PCR run producing two amplicons of 354 bp and 187 bp, respectively.

### Establishment of cFAK and FAKΔe4 overexpressing cell lines

For overexpression experiments, MDA-MB-231 cells were transduced with either canonical FAK (cFAK) or FAK isoform lacking exon 4 (FAKΔe4) with linker-eGFP sequence (on the C-terminus end) plasmids that were inserted into the pLenti4/V5-DEST vector (ThermoFisher). For plasmids, sequences from NCBI NM_001352694.2 and NM_001387652 were used for cFAK and FAKΔe4, respectively. These plasmids were created by Invitrogen GeneArt. Lentivirus particles were produced by transfecting each plasmid together with lentiviral expression vectors and third-generation packing constructs (GAG/POL, Rev and VSVG) in TransIT-LT1 (#MIR2304, Mirus Bio) into HEK293T cells (#CRL-3216, ATCC). Lentiviral particles were harvested and transduced into MDA-MB-231 cells in the presence of polybrene (#TR-1003, MilliporeSigma). After transduction, cells were sorted into different plasmid-expression cell populations using FACS. Overexpression efficiency was confirmed by western blotting.

### FACS-mediated sorting of transduced MDA-MB-231 cell populations

Prior to cell sorting, cells were kept in PBS-10% FBS and filtered using a 50 μm filter to remove biological fragments. After the transduction of the MDA-MB-231 cell line, modified MDA-MB-231 cells were sorted using the BD FACSAria™ Fusion flow cytometer to collect homogenous plasmid-expressing cell population series using eGFP expression: MDA-MB-231 cFAK^high^, MDA-MB-231 FAKΔe4^mid^ and MDA-MB-231 FAKΔe4^high^ cell lines. MDA-MB-231 wild-type cells were used as a baseline for autofluorescence.

### Polyacrylamide gels

PA gels of different stiffnesses were prepared as described before (84,85). Briefly, the ratio of acrylamide (40% w/v; #A0113, Bio-Rad) to bis-acrylamide (2% w/v; #A0114, Bio-Rad), in Milli-Q water and tetramethyl ethylenediamine (TEMED, #T9281, Sigma Aldrich) solution in 0.25 M HEPES buffer at pH 6, was adapted according to the range of desired stiffnesses from 0.5 to 50 kPa. Ammonium persulfate (APS, #BP179-100, Fisher Scientific) was used to initiate gel polymerization. The gels were functionalized using Sulfo-SANPAH (#PI22589, Fisher Scientific) under UV irradiation. Then, the PA gels were coated with 0.1 mg.mL^-1^ rat tail type I collagen (#354236, Corning). After sterilization with UV light, cells were seeded on top of the PA gels.

### Stiffness gradient polyacrylamide gel synthesis and durotaxis assay

Stiffness gradient PA gels were fabricated as previously described (86). Two acrylamide gel droplets of 1 and 30 kPa were added in the opposite side of a single activated coverslip. The 30 kPa acrylamide drop containing 0.5 µm red fluorescent microbeads (#F8812, Invitrogen) which, once merged with the 1 kPa solution, allows the production of a stiffness map due to the linear correlation between microsphere density and stiffness. A hydrophobic coverslip was gently dropped onto both solutions so that they would merge, resulting in a continuous stiffness gradient. To assess the optimal stiffness of MDA-MB-231 cell lines, 15,000 cells were seeded on collagen-coated stiffness gradient substrates and incubated for 72 hours as previously described (13). Cells were fixed with PBS-4% paraformaldehyde (PFA) for 15 minutes and nuclei were stained with DAPI for 10 minutes. The whole gel was imaged using the tile scan function of an Axio Observer.Z1/7 epifluorescence microscope equipped with an Axiocam 506 monochromatic camera using a x20/0.5 NA objective (Zeiss). Cell density was quantified within different stiffness regions using a custom MATLAB code.

### TCGA data analysis

The results analyzed here are based upon data generated by the The Cancer Genome Atlas (TCGA) Research Network (https://www.cancer.gov/tcga) from breast invasive carcinoma [TCGA-BRCA] containing primary tumor samples from 1,093 individual patients. Alternative splicing information in the form of specific exon read counts (GRCh37/hg19) of the FAK gene was obtained from the TCGA Splicing Variation Database (http://www.tsvdb.com/index.html). Additional clinical data and RNA-seq read counts for gene expression were downloaded from the TCGA database. Patient samples with low KRT8 gene expression (<10,000 reads) corresponding to stroma-biased samples were excluded. FAK exon 4 percent spliced-in (PSI) values were calculated based on the ratio of FAK exon 4 inclusion events (chr8:141874498-141889570 and chr8:141889736-141900642) and FAK exon 4 exclusion events (chr8:141874498-141900642). The YAP signature was produced using a score of 14 co-transcription target genes of YAP1 to rank the patient samples as described before (30). The same strategy was used for colon adenocarcinoma (COAD), cholangiocarcinoma (CHOL), lung squamous cell carcinoma (LUSC), stomach adenocarcinoma (STAD), rectum adenocarcinoma (READ) and esophageal carcinoma (ESCA) TCGA datasets. For survival curves, data were divided into a low and high FAKΔe4 level groups based on an expression threshold computed in R with the surv_cutpoint function (survminer v0.5.2). These two groups were then used to compute the Kaplan-Meier curves and associated p-values in R using the survfit and ggsurvplot functions.

### Library preparation, long-read RNA-sequencing and data processing

Cell lysates were homogenized with QIAshredder columns (#79654, QIAGEN,) and total RNA was extracted with RNeasy mini columns (#74104, QIAGEN,). RNA was DNase-treated using TURBO DNase (#AM2238, Thermo Fisher Scientific,). RNA quality was verified on a 2100 Bioanalyzer (Agilent Technologies) and polyadenylated RNA was enriched using the NEBNext Poly(A) mRNA Magnetic Isolation Module (#E7490L, New England Biolabs). Libraries were generated using 100 ng of poly(A)+ RNA and Oxford Nanopore Technologies (ONT)’ Direct cDNA Sequencing Kit (#SQK-DCS109, Oxford Nanopore Technologies), with each sample barcoded with the Native Barcoding Expansion (#EXP-NBD104, ONT), according to the manufacturer’s specifications. Barcoded samples were pooled, loaded on a MinION sequencer using four R9 Flow Cells (#FLO-MIN106D, ONT), and sequenced using the MinKNOW software v19. Raw fast5 files were basecalled using Guppy v.4.2.2 with the guppy_basecaller command and the dna_r9.4.1_450bps_fast.cfg configuration file and default settings. Barcodes were detected and reads were separated with guppy_barcoder using the default settings. The resulting FASTQ reads were aligned to the GRCh38 human reference with Minimap2 v2.17-r941 (87) using the following parameters: -aLx splice –cs = long. Alternative splicing events were quantified using PSI-Sigma (v2.1), a junction-based approach for estimating PSI values from RNA-seq data. PSI-Sigma was executed via the dummyai.pl script with a minimum read support threshold of five reads per event (--nread 5). Splicing analyses were performed using the same Ensembl genome annotation.

### RNA sequencing public data retrieval and data processing

mRNA sequencing (RNA-seq) data were obtained from the Gene Expression Omnibus (GEO) under accession numbers GSE205816 (MCF10CA1a cell line) and GSE255829 (MDA-MB-231 cell line). Raw sequencing reads (FASTQ format) were processed using fastp (v0.23.4) for quality control and adapter trimming with default parameters, including automatic detection of paired-end adapters, trimming of bases with Phred quality scores < 20, removal of reads containing > 30% low-quality bases, and exclusion of reads shorter than 30 bp. Once raw files from public data and long-read RNA sequencing were obtained, high-quality reads were aligned to the human reference genome (GRCh38) using STAR (v2.7.3a) with default settings, and resulting alignments were retained as coordinate-sorted BAM files. Alternative splicing events were quantified using PSI-Sigma (v2.1). PSI-Sigma was executed via the dummyai.pl script with a minimum read support threshold of five reads per event (--nread 5). Splicing analyses were performed using the same Ensembl genome annotation.

### Western blotting

Live cells on PA gels were directly lysed using preheated Laemmli buffer. Total protein extracts were stored at -80°C until use. For electrophoresis, protein samples underwent a 9% acrylamide gel run and western blotting. Membranes were blocked with 5% milk in 0.1% tween-TBS, or 5% BSA in 0.1% tween-TBS for the detection of phosphorylated proteins. Membranes were incubated overnight with the primary antibody at 4 °C and 1 hour with the secondary antibody at room temperature. Total FAK and pY397-FAK primary antibodies were prepared at 1:5,000 dilution, GAPDH primary antibody at 1:20,000 dilution and the secondary antibody was diluted at 1:5,000. Membranes were imaged with the West Pico PLUS chemiluminescent substrate (#PI34577, Thermo Fisher Scientific) using the ImageQuant 800 system (Cytiva). Quantifications were performed with ImageJ.

### Immunofluorescence on fixed samples

At least 4×10^4^ cells were seeded on PA gels or glass for an overnight incubation to allow for adequate attachment and spreading. Cells were fixed for 10 minutes with PBS-4% PFA. Cells were permeabilized and blocked for 1 hour in PBS containing 5% goat serum and 0.1% Triton X-100 (#9002-93-1, MilliporeSigma). The primary antibody was incubated in PBS-1% goat serum for 2 hours. The secondary antibody was incubated in PBS-1% goat serum for 1 hour. The nuclei were stained with DAPI (#D1306, Thermo Fisher Scientific) for 10 minutes. Then, samples were mounted onto larger glass slides (#5075-W, Brain Research Laboratories) using Fluoromount-G (#0100-01, Cedarlane). The coverslip edges were sealed with nail polish. The slides were stored in the dark at 4 °C until imaging.

### Focal adhesion timelapse

To assess FA dynamics, 8×10^5^ transduced MDA-MB-231 cells were seeded in a glass bottom fluorodish (#FD35-100, World Precision Instruments) for an overnight incubation. A Nikon H-TIRF Ti microscope was used with a 60X oil objective, 70-80 μm depth and captured with an ORCA-ERA camera. FAs were captured every 120 seconds for 80 minutes. Image stacks were analyzed using the “multi measure” feature of ImageJ of selected FA regions of interest. Color-coded projections were generated using LUT “Physics”.

### FRAP experiments

For FRAP experiments, 8×10^5^ transduced MDA-MB-231 cells were seeded in a glass bottom fluorodish (#FD35-100, World Precision Instruments) the day before. A LSM 900 confocal microscope (Zeiss) was used with the Plan-Apochromat 40X/1.4 Oil DIC objective. The FA region of interest was first selected, followed by the acquisition of 10 images before photobleaching to quantify the baseline for the recovery. After using the 488 nm laser at 100% with 10 iterations for area bleaching, 1 image every 1 second was captured for 60 seconds. Of note, only one focal adhesion (FA) was photobleached per cell. Image stacks were analyzed using the “multi measure” feature of ImageJ of selected FA regions of interest to quantify the recovery rate of fluorescence signal.

### 2D migration assay

PA gels of 1, 5, 10, 18, 30 and 50 kPa were fabricated, placed in a 6-well plate and fixed with vacuum grease. Once the plate was sterilized with UV light, 1×10^5^ transduced MDA-MB-231 cells were seeded for an overnight incubation. The Axio Observer.Z1/7 microscope (Zeiss) was used with an Axiocam 506 monochromatic camera for the brightfield capture and an Axiocam 705 color camera for fluorescence capture, with a x5/0.13 NA. For the acquisition settings, 10 positions per condition were chosen and were first captured with the 488 nm fluorescence to ensure that analyzed cells were eGFP-positive. Then, 1 image every 15 minutes was captured for 12 hours with brightfield illumination. To prevent the loss of tracked cells that went off-screen along the experiment, only 6 hours of the 12 hours migration were analyzed. To do so, the manual tracking feature of ImageJ was used to quantify cell migration.

### Collagen hydrogel preparation

The collagen was isolated from rat tails as previously described (83). The collagen was extracted from rat tail tendons and solubilized in 0.1% acetic acid to make a 10 mg.mL^-1^ stock solution. On ice, hydrogels were prepared by diluting the stock solution with 0.1% acetic acid and neutralized with a 10X D-PBS containing 25 mM HEPES, 44 mM sodium bicarbonate and 0.11 g.L^-1^ phenol red addition with a 1 M NaOH solution. For glycated collagen, collagen stock solutions (10 mg.mL ¹ in 0.1% sterile acetic acid) were mixed with 500 mM ribose stock solutions. The resulting mixtures contained either 0 or 100 mM ribose and were incubated for 5 days at 4 °C. After glycation, 10× PBS-HEPES and NaOH were added as described above to obtain a final collagen concentration of 1.5 mg.mL ¹ before casting in the cantilevers as previously described (88).

### Spheroid invasion assay

The cell suspension containing 5×10^4^ cells.mL^-1^ was distributed in an ultra-low attachment 96 microwell plate (#83.3925.400, Sardstedt) at 100 µL per well in the MCF10a medium. The plate was incubated for 4 days to allow optimal spheroid formation where the medium was changed every 48 hours. The spheroids were then added into the collagen solution, of which 30 µL was seeded in a glass bottom 24-well plate (#P24G-1.5-13-F, MatTeK) and inverted at least thrice over 30 minutes while being maintained at 37 °C. Subsequently, 200 µL of collagen solution were added and the mixture was allowed to polymerize at 37 °C for 1 hour before adding 500 µL of DMEM. The medium was changed thrice over 90 minutes for both glycated (100 mM ribose) and non-glycated (0 mM ribose) collagen hydrogels once polymerized to remove the excess of ribose from the hydrogel. Of note, 1.5 mg/mL collagen gels with 0 or 100 mM ribose form gels have comparable fiber architecture but tunable stiffness (47). The embedded spheroids were allowed to invade for 3 days. The medium was changed every 48 hours. Brightfield images were taken on days 0, 1, 2 and 3 using an Axio Observer.Z1/7 epifluorescence microscope equipped with an Axiocam 506 monochromatic camera using a x5/0.13 NA objective (Zeiss).

### Migration wound healing assay

Wells of a 96-well plate (#35072, Falcon) were coated with 50 µL of collagen type I (#354236, Corning) at 300 µg.mL^-1^ (diluted in HEPES at 50 mM, pH 8) for 2 hours at 4 °C. After coating solution removal, 100 µL of PBS were added, followed by UV irradiation for 1 hour. For both cells lines (MDA-MB-231 cFAK^high^ and FAKΔe4^high^), 50,000 cells per well were seeded with 100 µL of final volume in each well and incubated overnight in an incubator at 37°C with 5% CO_2_ overnight. The day after, when cells reached full confluence, wounds were created using The Incucyte® 96-Well Woundmaker Tool (#4563, Sartorius) (3 runs to ensure the proper creation of the wound). Two washes were performed with 100 µL of medium. Acquisitions were performed using a Sartorius Incucyte® SX5 in Scratch Wound mode with a 10X objective every 2 hours for 48 hours. The Incucyte® Scratch Wound Analysis (#9600-0012, Sartorius) started after 30 minutes to allow the plate to warm to 37 °C. The analysis was done on ImageJ using the previously published macro “Wound_healing_size_tool_updated.ijm” (89).

### Statistical analysis

All graphs and statistical analysis were performed using PRISM (GraphPad). For normally distributed data, the statistical significance was computed using one-way ANOVA or two-way ANOVA with Tukey’s correction for multiple comparisons or student’s *t*-test unless specified otherwise. The statistical significance was also computed using a non-parametric Kruskal-Wallis test with Dunn’s correction for multiple comparisons.

