## Supplementary figures for "Matrix Stiffness-driven FAK Splicing Tunes Cell Mechanosensing"

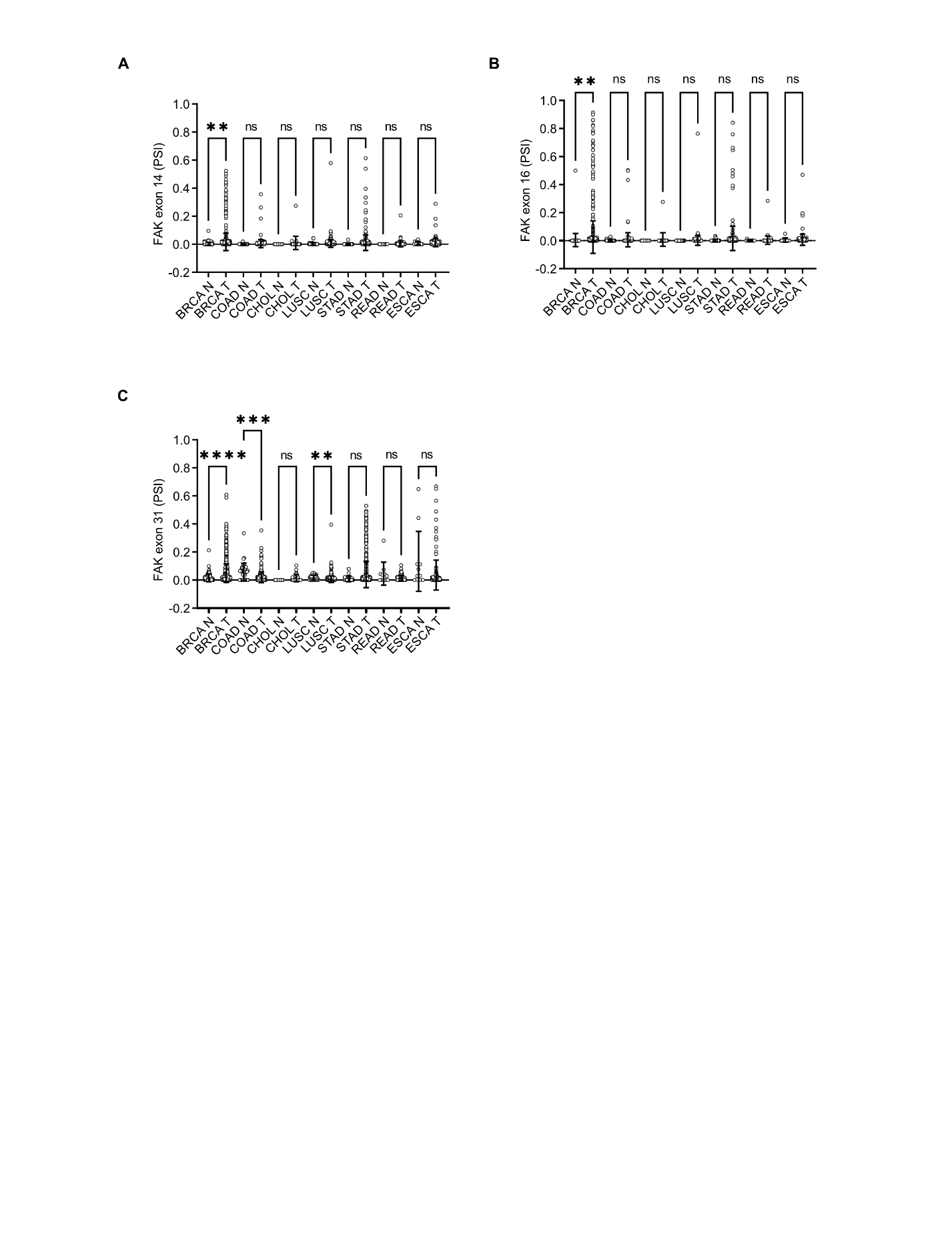
**Supplementary Figures**

**Fig.S1. Alternative splicing profiles of FAK exons 14, 16 and 31 in patients.** (**A**) Percent spliced-in (PSI) of FAK exon 14, (**B**) exon 16 and (**C**) exon 31 in solid tumor tissues (T) compared to the corresponding normal tissues (N) in breast cancer (BRCA), colon adenocarcinoma (COAD), cholangiocarcinoma (CHOL), lung squamous cell carcinoma (LUSC), stomach adenocarcinoma (STAD), rectum adenocarcinoma (READ) and esophageal carcinoma (ESCA) using TCGA datasets. One-way ANOVA with Tukey. Data are means ± SEM; **p<0.01, ***p<0.001, ****p<0.0001.

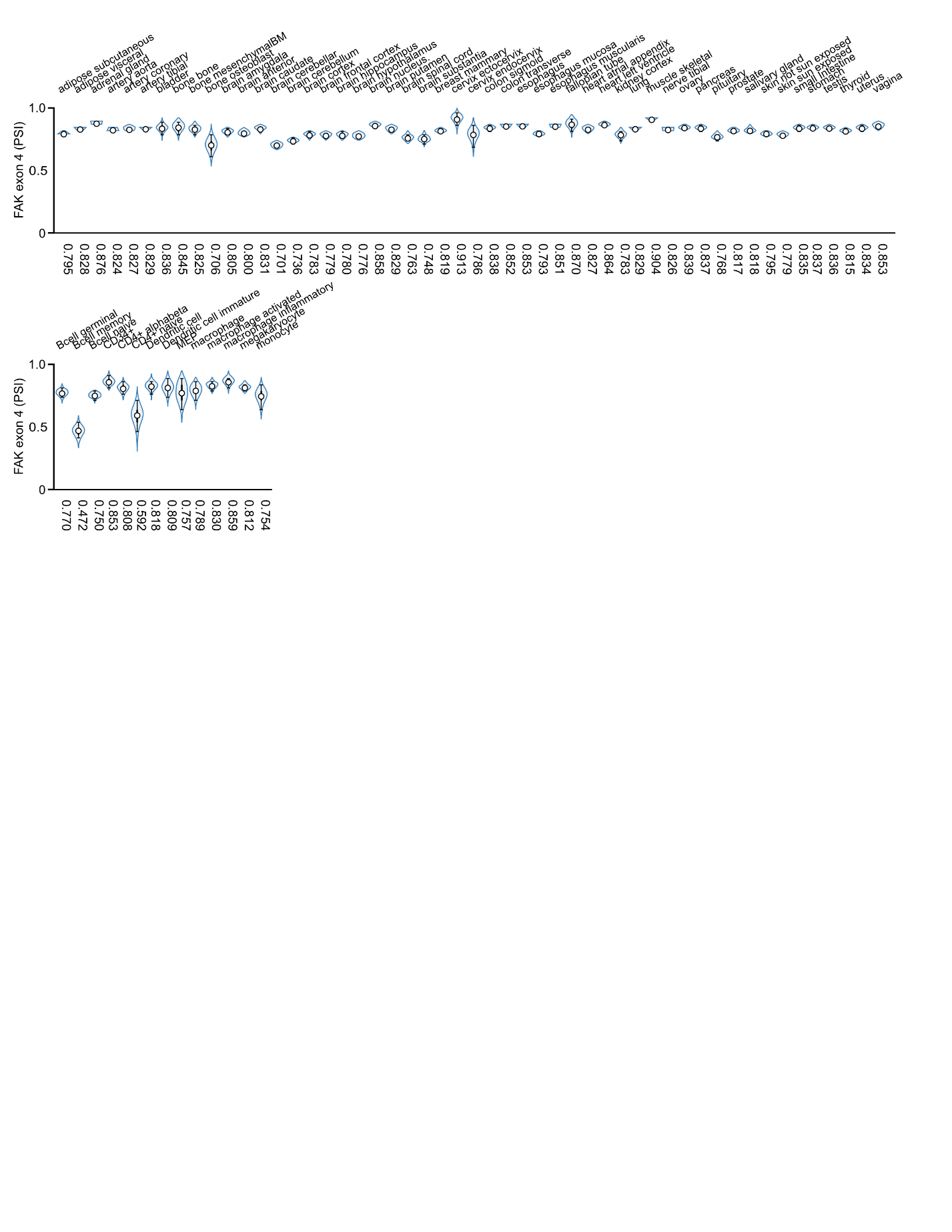

**Fig.S2. FAK∆e4 is ubiquitous.** Graphical output of MAJIQlopedia showing average PSI of FAK exon 4 across healthy tissues.

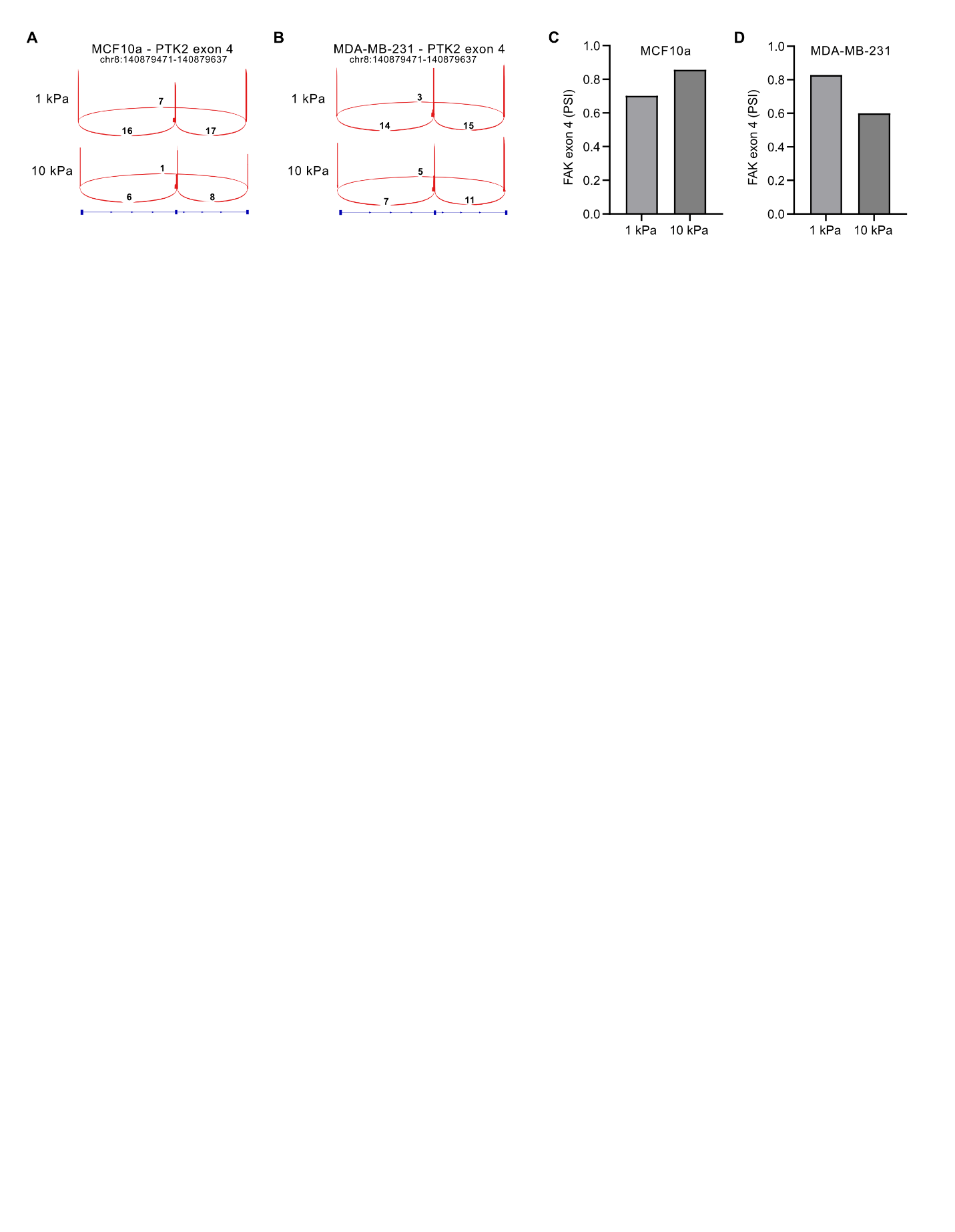
**Fig.S3. FAK exon 4 PSI of MCF10a and MDA-MB-231 cells on substrates of varying stiffness using long-read RNA sequencing.** (**A**) Sashimi plots showing inclusion and exclusion events of FAK exon 4 in MCF10a and (**B**) MDA-MB-231 cells cultured on 1 and 10 kPa. N=1. (**C**) Corresponding quantification of FAK exon 4 PSI in MCF10a and (**D**) MDA-MB-231 cells cultured on 1 and 10 kPa.

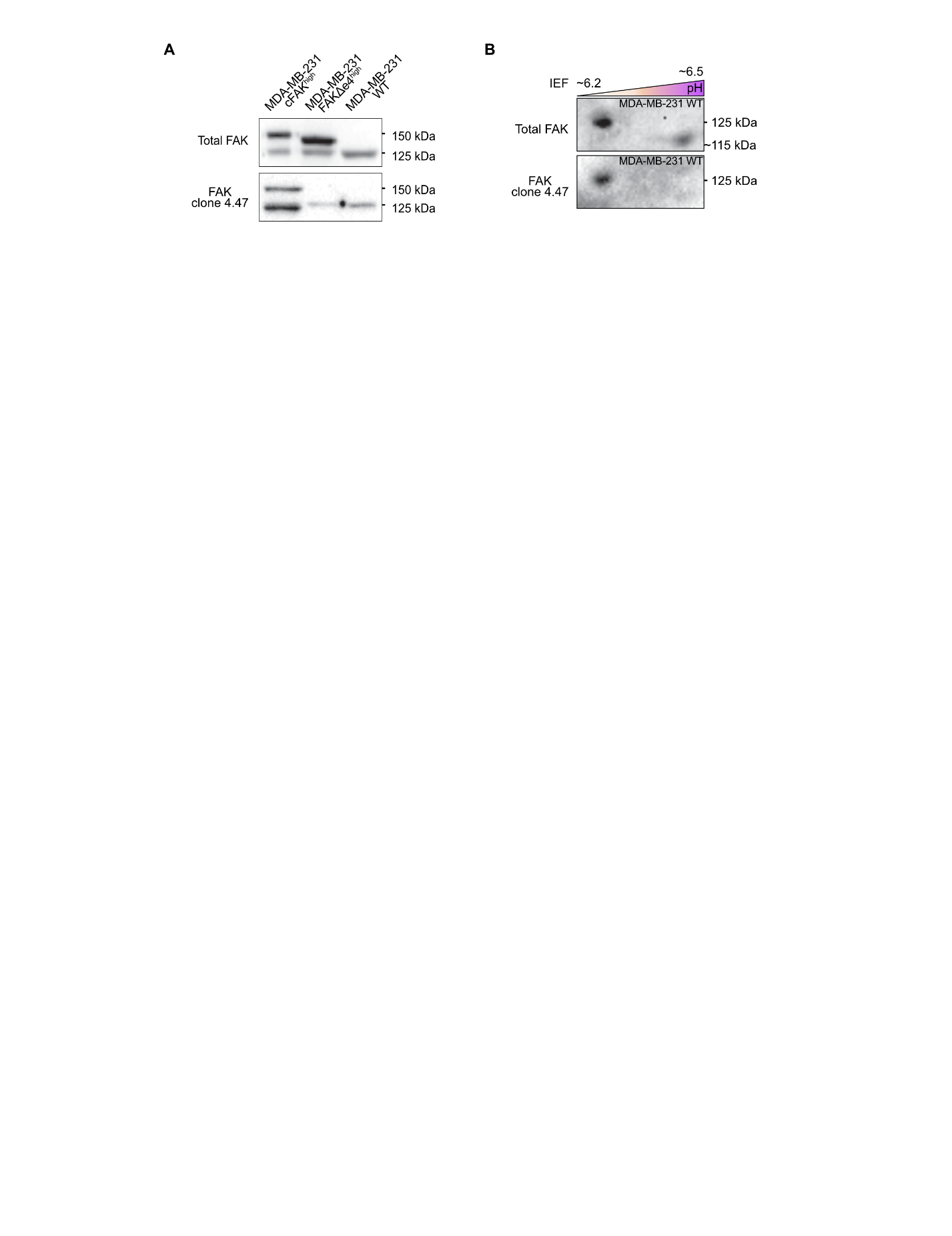

**Fig.S4. FAK∆e4** **is mechanoregulated at protein level.** (**A**) Western blots of whole protein extracts from MDA-MB-231 cFAK^high^, FAK∆e4^high^ and WT showing total FAK and FAK clone 4.47 chemiluminescent signals. (**B**) 2D western blots of a whole protein extract from MDA-MB-231 cells (WT), showing pH gradient on the x axis and molecular weight gradient on the y axis performed using total FAK and FAK clone 4.47 antibodies.

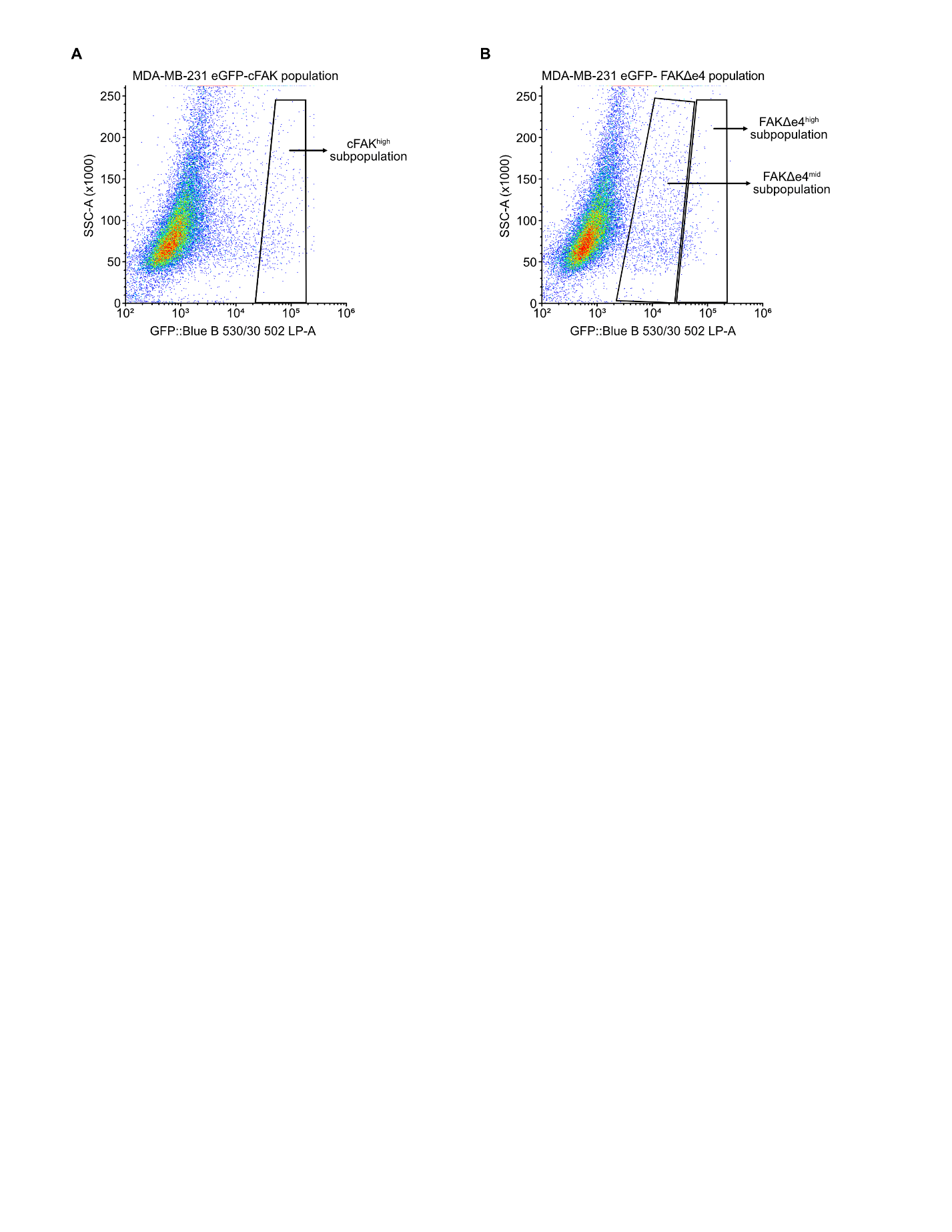

**Fig.S5. Cell sorting of MDA-MB-231 eGFP-cFAK and eGFP-FAK∆e4 subpopulations.** (**A**) Dot plots of the distribution of MDA-MB-231 eGFP-cFAK and (**B**) MDA-MB-231 eGFP-FAK∆e4 cell populations based on the eGFP expression intensity. Cut-offs for the selection of the subpopulations (MDA-MB-231 cFAK^high^, FAK∆e4^mid^ and FAK∆e4^high^) are shown.

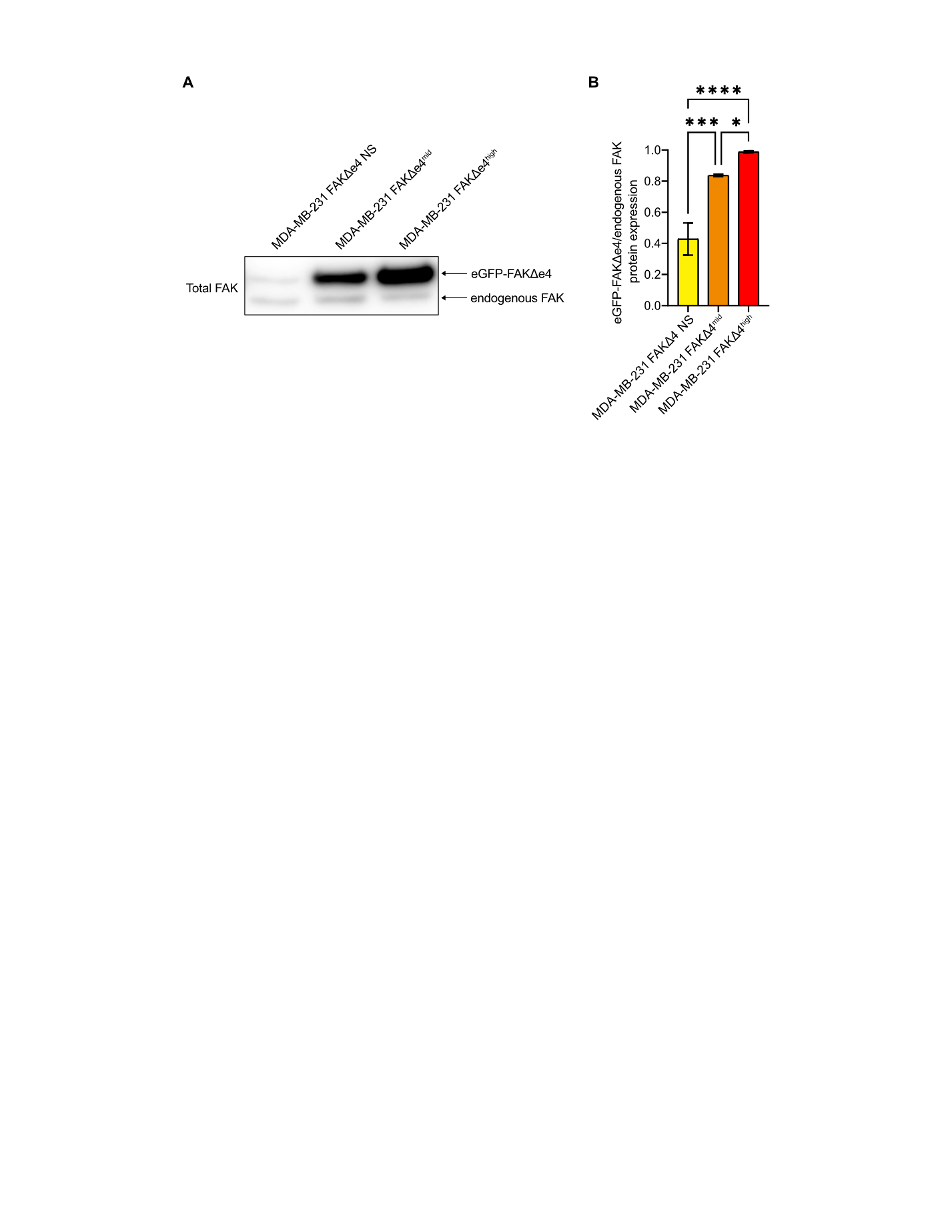

**Fig.S6. MDA-MB-231 cell populations with FAK∆e4 overexpression.** (**A**) Western blot of total FAK using whole protein extracts from MDA-MB-231 FAK∆e4 non-sorted (NS), FAK∆e4^mid^ and FAK∆e4^high^ cells cultured on glass for 48 hours. (**B**) Corresponding densitometric quantification of exogenous FAK∆e4/endogenous FAK ratios for MDA-MB-231 FAK∆e4 NS, FAK∆e4^mid^ and FAK∆e4^high^ cells. N=3. One-way ANOVA with Tukey. Data are means ± SEM; *p<0.05, ***p<0.001, ****p<0.0001.

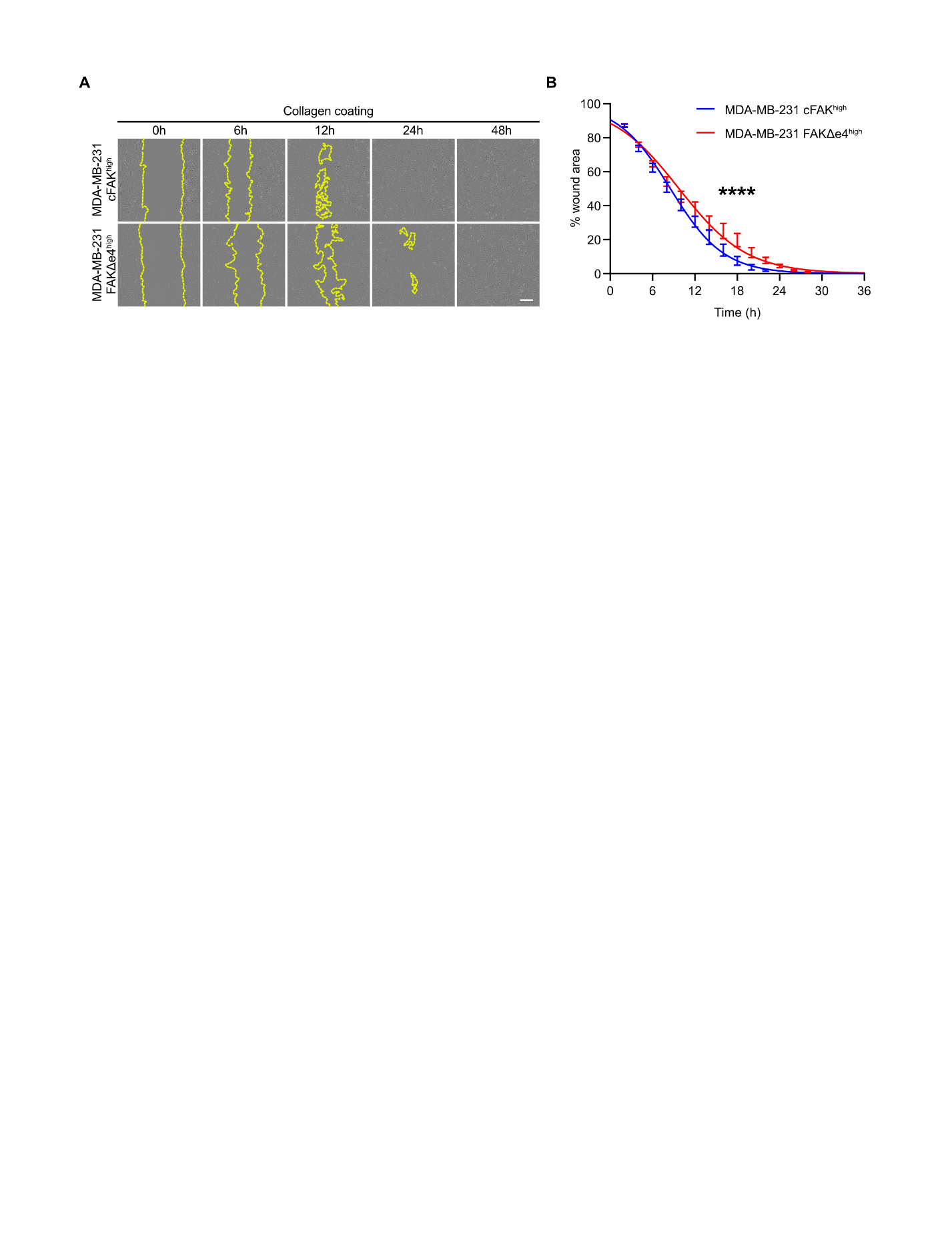
**Fig.S7. Collective cell migration.** (**A**) Representative images of cell migration of MDA-MB-231 cFAK^high^ and FAK∆e4^high^ cells on collagen-coated plastic after 0, 6, 12, 24 or 48 hours using a wound healing assay. (**B**) Corresponding quantification of the wound closure over time by MDA-MB-231 cFAK^high^ and FAK∆e4^high^ cells. N=3. Extra sum-of-squares F test with LogEC50 and Hill Slope parameters. Data are means ± SEM.; ****p<0.0001. Scale bar = 200 μm.

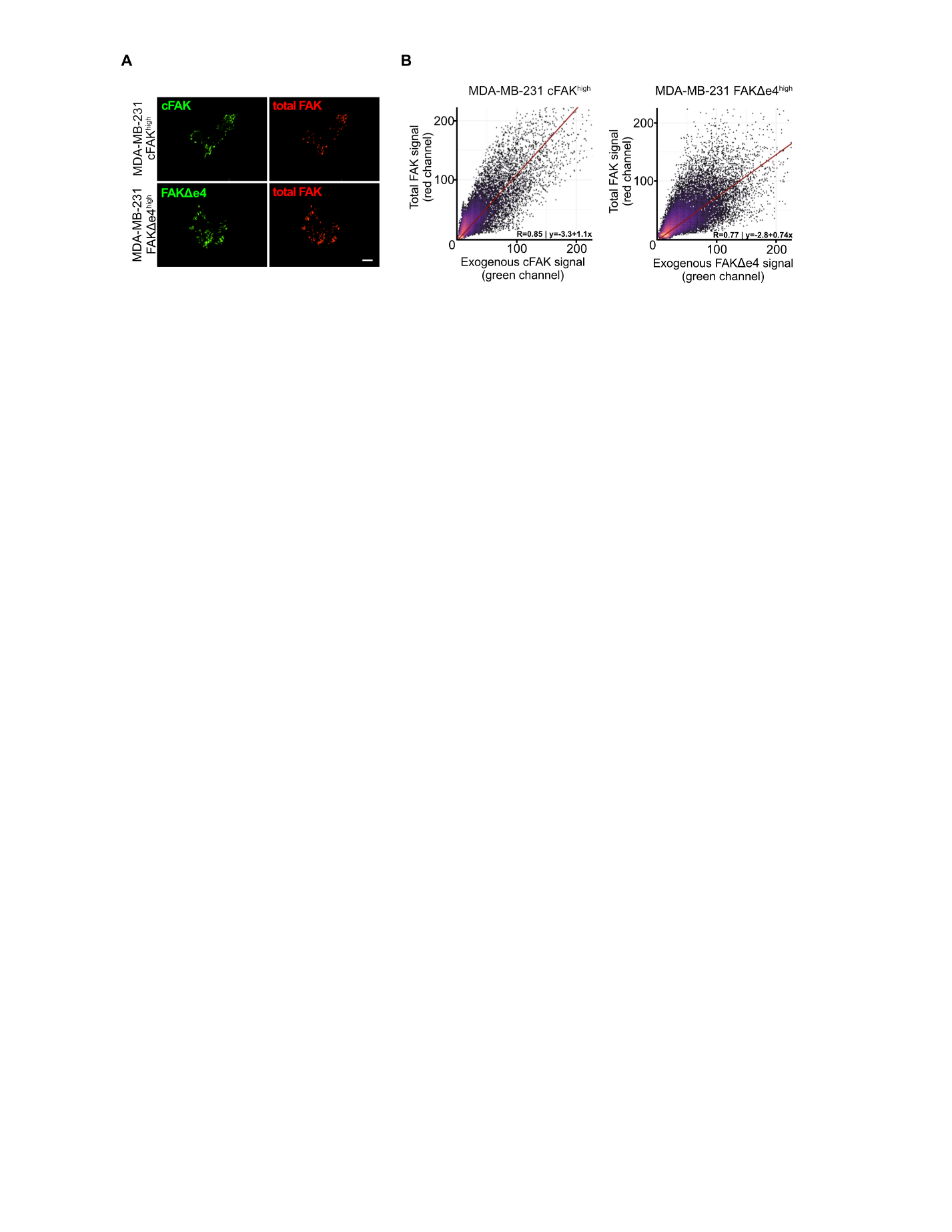

**Fig.S8. Colocalization of FAK∆e4 with total FAK at focal adhesions.** (**A**) Representative immunofluorescence of MDA-MB-231 cFAK^high^ and FAK∆e4^high^ cells with eGFP staining for exogenous FAK (green) and total FAK (red) using TIRF microscopy at FAs. Scale bar = 10 μm. (**B**) Representative cytofluorograms comparing pixel-by-pixel showing colocalization of total FAK vs. exogenous FAK (cFAK or FAK∆e4).

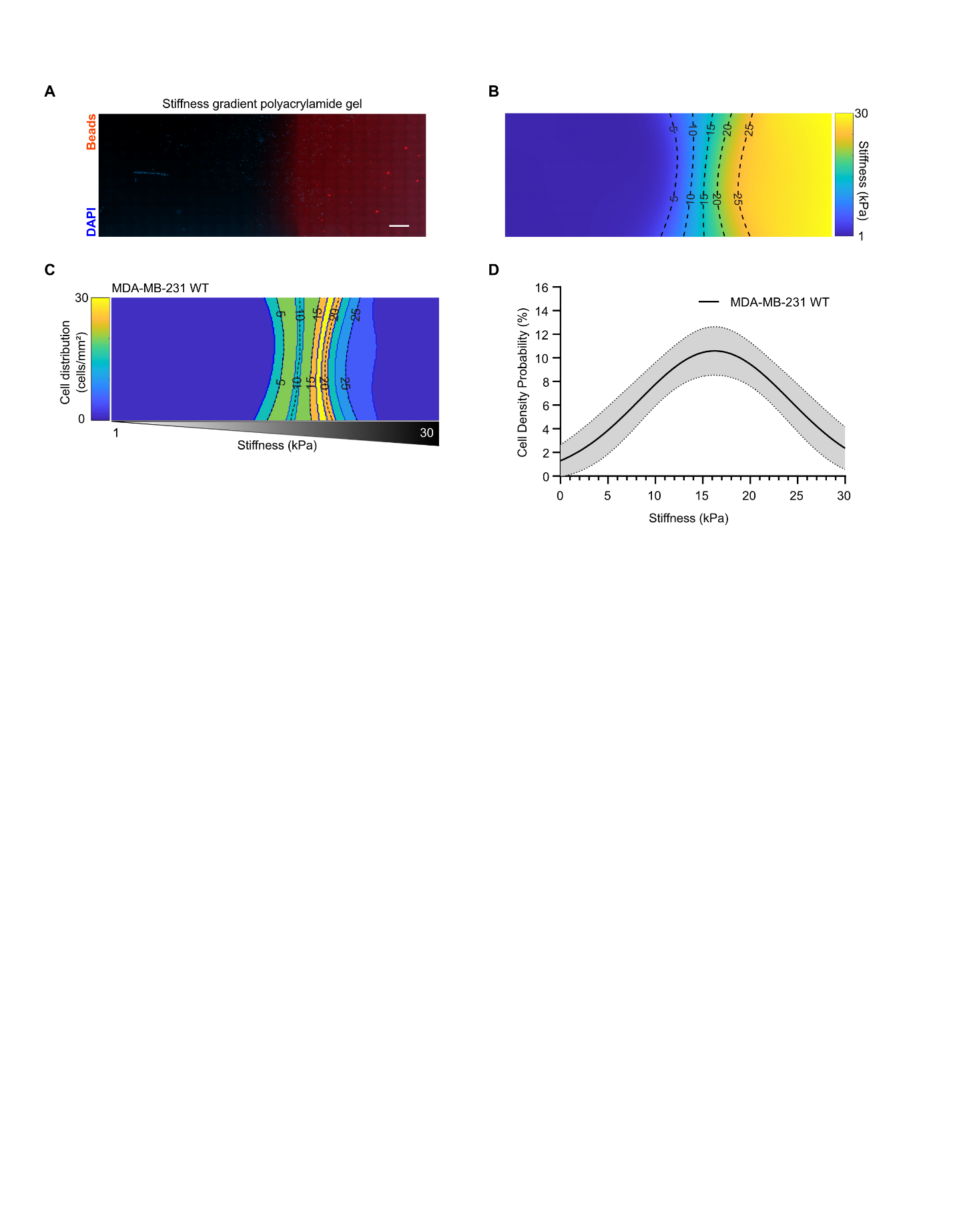

**Fig.S9. Durotaxis of parental MDA-MB-231 cells.** (**A**) Tile scan image of a stiffness gradient PA gel ranging from 1 to 30 kPa. Red fluorescent beads were used to determine the stiffness gradient. Cells were stained with DAPI. Scale bar = 150 μm. (**B**) Corresponding stiffness gradient determined using a MATLAB script. (**C**) Representative heat map of cell density probability as a function of substrate stiffness from 1 to 30 kPa for MDA-MB-231 WT cells. (**D**) Corresponding quantification of the cell density probability according to substrate stiffness presented as a Gaussian fit with 95% confidence interval. N=3.
